# Impairment of SARS-CoV-2 spike-glycoprotein maturation and fusion-activity by nitazoxanide: an effect independent of spike variants emergence

**DOI:** 10.1101/2021.04.12.439201

**Authors:** Anna Riccio, Silvia Santopolo, Antonio Rossi, Sara Piacentini, Jean-Francois Rossignol, M. Gabriella Santoro

## Abstract

SARS-CoV-2, the causative agent of COVID-19, has caused an unprecedented global health crisis. The SARS-CoV-2 spike, a surface-anchored trimeric class-I fusion-glycoprotein essential for viral entry, represents a key target for developing vaccines and therapeutics capable of blocking virus invasion. The emergence of SARS-CoV-2 spike-variants that facilitate virus spread and may affect vaccine efficacy highlights the need to identify novel antiviral strategies for COVID-19 therapy. Here we demonstrate that nitazoxanide, an antiprotozoal agent with recognized broad-spectrum antiviral activity, interferes with SARS-CoV-2 spike biogenesis, hampering its maturation at an endoglycosidase H-sensitive stage. Engineering multiple SARS-CoV-2 variant-pseudoviruses and utilizing quantitative cell-cell fusion assays, we show that nitazoxanide-induced spike modifications hinder progeny virion infectivity as well as spike-driven pulmonary cell-cell fusion, a critical feature of COVID-19 pathology. Nitazoxanide, being equally effective against the ancestral SARS-CoV-2 Wuhan-spike and different emerging variants, including the Delta variant of concern, may represent a useful tool in the fight against COVID-19 infections.

## INTRODUCTION

Human coronaviruses (HCoV) were discovered in the 1960s and were originally thought to cause only mild upper respiratory tract diseases in immunocompetent hosts, with several strains being responsible for the common cold [1–3]; however, public awareness of HCoV has changed considerably since the beginning of this century, with the SARS (Severe Acute Respiratory Syndrome) epidemic in 2002 and the MERS (Middle East Respiratory Syndrome) outbreak in 2012, two zoonotic CoV infections that resulted in mortality rates of approximately 10% and 35%, respectively [4].

Near the end of 2019 the seventh human coronavirus, phylogenetically in the SARS-CoV clade (lineage B) and therefore named SARS-CoV-2, emerged in China [5]. SARS-CoV-2 turned out to be a far more serious threat to public health than SARS and MERS HCoVs because of its ability to spread more efficiently than its predecessors, making it difficult to contain worldwide, and causing an unprecedented global health crisis, as well as societal and economic disruption. The clinical features of the disease associated with SARS-CoV-2 infection, COVID-19 (coronavirus disease-2019), vary ranging from asymptomatic state to respiratory symptoms that, in a subset of patients, may progress to pneumonia with massive alveolar damage and extreme rise in inflammatory cytokines production that may lead to acute respiratory distress syndrome (ARDS), thrombosis, multi organ dysfunction and death [5, 6].

SARS-CoV-2 infection is initiated by the binding of the spike (S) glycoprotein to the host receptor, which, similarly to SARS-CoV, is the human angiotensin-converting enzyme 2 (hACE2)[7–9]. The spike protein, together with the M (membrane) and E (envelope) proteins, is anchored into the viral envelope, decorating the virion surface as a distinctive crown (“corona”) (Fig. 1A), and is essential for viral entry into target cells [7, 8].

**Figure 1.**
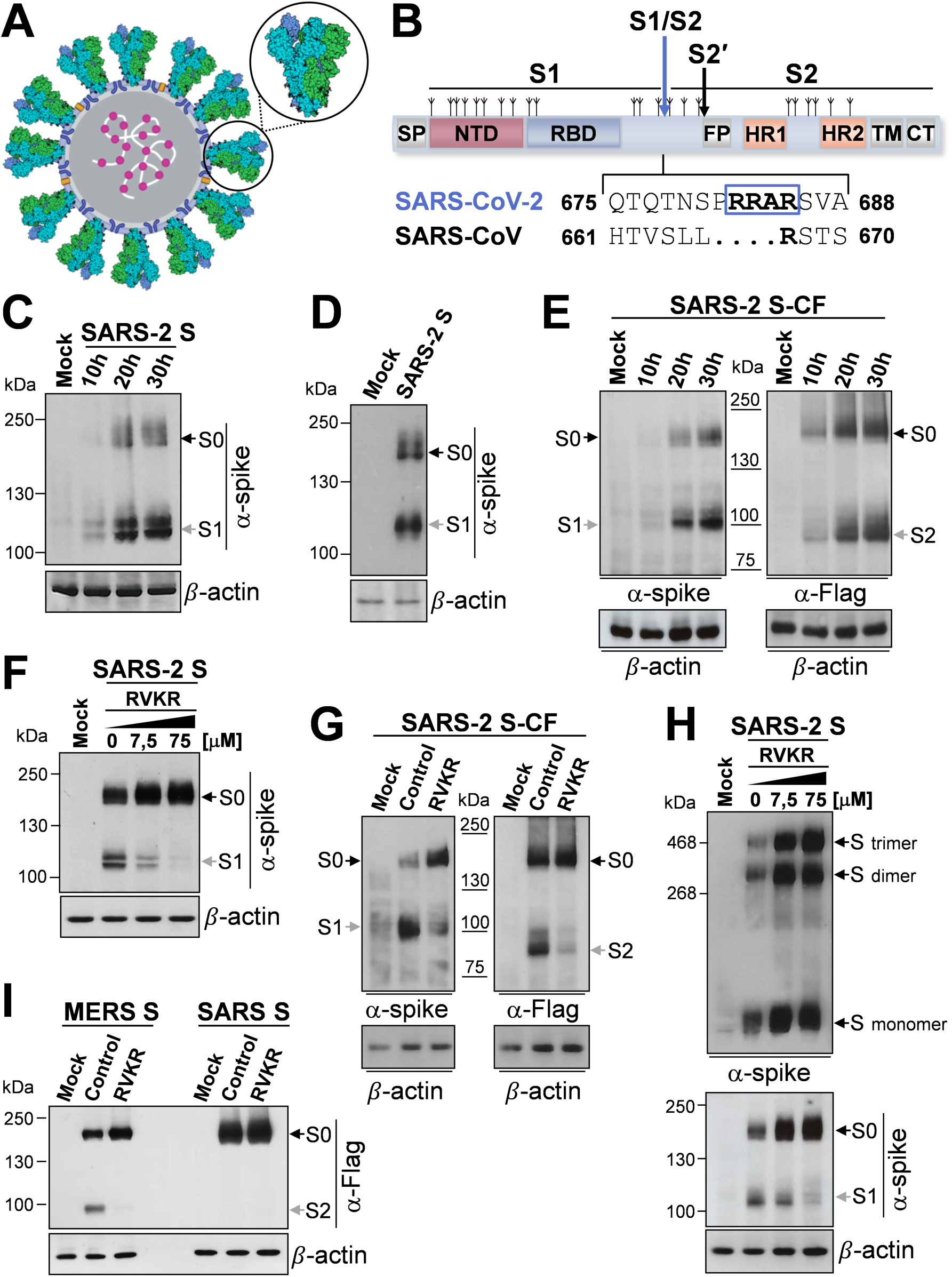
Characterization of SARS-CoV-2 spike biogenesis, furin-mediated S1/S2 cleavage and trimerization in human lung cells. (**A**) Schematic representation of the SARS-CoV-2 virus structure. The lipid bilayer comprising the spike glycoprotein S (indicated by the circle), the membrane protein M (blue) and the envelope protein E (yellow), and the viral RNA (white) associated with the nucleocapsid protein N (magenta) are shown. 3D-model of the spike glycoprotein homotrimer (PDB ID: 6VSB) is shown in the zoom; each spike monomer is colored individually. (**B**) Schematic illustration of SARS-CoV-2 S glycoprotein. S1, receptor-binding subunit; S2, membrane fusion subunit. The positions of N-linked glycosylation sequons are shown as branches. Protein domains are illustrated: SP, signal peptide; NTD, N-terminal domain; RBD, receptor binding domain; FP, fusion peptide; HR1, heptad repeat 1; HR2, heptad repeat 2; TM, transmembrane domain; CT, cytoplasmic tail. The S1/S2 and S2′ protease cleavage sites are indicated by arrows. Sequence comparison of the S1/S2 cleavage site of SARS-CoV and SARS-CoV-2 spike glycoproteins, and the putative furin cleavage site (RRAR residues in the box) in SARS-CoV-2 are shown. GenBank accession numbers are AFR58740.1 for SARS-CoV S and QHD43416.1 for SARS-CoV-2 S [85]. (**C,D**) Detection of SARS-CoV-2 S protein (α-spike) levels by immunoblot (IB) in whole cell extracts (WCE) from human lung A549 epithelial cells (**C**) or MRC-5 fibroblasts (**D**) transiently transfected with the SARS-CoV-2 spike construct (SARS-2 S) or empty vector (Mock) at different times (**C**) or at 20h (**D**) after transfection. (**E**) A549 cells were transiently transfected with the C-terminal Flag-tagged SARS-CoV-2 spike construct (SARS-2 S-CF) or empty vector and, at different times, WCE were analyzed for S protein levels by IB using anti-spike or anti-Flag antibodies. (**F,G**) Levels of S protein determined by IB using anti-spike or anti-Flag antibodies in WCE from A549 cells transiently transfected with the SARS-2 S (**F**) or SARS-2 S-CF (**G**) constructs, or empty vector for 4h and treated with different concentrations (**F**) or 50 μM (**G**) of furin inhibitor decanoyl-RVKR-CMK (RVKR) or vehicle (Control) for 16h. (**H**) Oligomeric status of SARS-CoV-2 spike protein in A549 cells. Gel electrophoresis (4% polyacrylamide) of WCE from A549 cells transfected with SARS-2 S construct or empty vector for 4h and treated with different concentrations of RVKR or vehicle for 16h. The different forms of the S protein were visualized by IB with the anti-spike antibody (top). Dimers and trimers are indicated. In parallel, S protein and S1 subunit levels in the same samples (8% polyacrylamide gels) are shown (bottom). (**I**) A549 cells were transfected with C-terminal Flag-tagged MERS-CoV spike (MERS S) or SARS-CoV spike (SARS S) constructs or empty vector and, after 4h, were treated with RVKR (50 μM) or vehicle (Control). After 16h, WCE were analyzed for levels of SARS-CoV or MERS-CoV S proteins by IB using anti-Flag antibodies. (**C-I**) Black arrows indicate bands corresponding to uncleaved S proteins (S0), whereas gray arrows indicate bands corresponding to the S1 or S2 subunits.

Similarly to other viral fusion glycoproteins, including influenza hemagglutinin (HA) and parainfluenza fusion (F) proteins, the SARS-CoV-2 spike is a trimeric class-I fusion protein; each monomer is synthesized as a fusogenically-inactive precursor of about 180 kDa containing an N-terminal signal peptide (SP) (Fig. 1B) that primes the nascent polyprotein for import into the endoplasmic reticulum (ER), where the protein is extensively modified with N-linked glycans (each protomer comprises 22 N-linked glycosylation sequons) [10, 11]. The spike protein assembles into an inactive homotrimer, which is endoproteolytically cleaved by cellular proteases giving rise to a metastable complex of two functional subunits: S1 (bulb) containing the receptor-binding domain (RBD) responsible for recognition and attachment to the host hACE2 receptor, and the membrane-anchored S2 (stalk) that contains the fusion machinery (Fig. 1B). S2 harbors the fusion peptide (FP), a short segment of 15–20 conserved mainly hydrophobic amino acids, which anchors to target membranes and plays an essential role in mediating membrane fusion by disrupting and connecting lipid bilayers of host cells. The FP is followed by two heptapeptide repeat sequences HR1 and HR2, the transmembrane anchor domain (TM), and a short cytoplasmic tail (CT) (Fig. 1B) [9].

S glycoproteins passing the quality control mechanisms of the ER are transported to the ER/Golgi intermediate compartment (ERGIC), the presumed site of viral budding [12]. Glycosylation plays an essential role in establishing viral spike proteins bioactive conformation and stability, for shaping viral tropism and has effects on virus antigenic properties, receptor binding and fusion activity [10,13,14].

Triggering of SARS-CoV-2 S fusion activity is a multistep process that requires sequential cleavage at distinct sites, including, in addition to the canonical CoV cleavage site at the S1/S2 boundary, an S2’ site located immediately upstream of the fusion peptide (Fig. 1B). In the case of SARS-CoV-2 an unexpected polybasic furin-like cleavage site (FCS), harboring multiple arginine residues, located at the S1/S2 boundary was shown to be processed during S protein biogenesis for S-protein “priming” [7,15,16]. Similar cleavage sites have been found in MERS-CoV, but not in other lineage B βCoVs, including SARS-CoV; importantly, the acquisition of similar cleavage sites is associated with increased pathogenicity in other viruses such as avian influenza viruses [17], and it is suggested to be responsible for SARS-CoV-2 high infectivity and transmissibility [7, 15].

The SARS-CoV-2 spike glycoprotein is also the primary target of host immune defenses and it is therefore the focus of vaccine development [18, 19]. At present, in parallel with a major vaccination endeavor, widespread whole genome sequencing efforts have allowed researchers to track the spread of different lineages globally, identifying several SARS-CoV-2 spike mutations that, providing fitness advantages, facilitate a rapid spread of the virus, also creating great concern on whether some variants may affect the efficacy of recently developed vaccines [20–23].

Therefore, in addition to prophylaxis, antiviral therapeutics remain urgently needed to combat SARS-CoV-2 in the current pandemic.

Nitazoxanide (NTZ), a thiazolide originally developed as an antiprotozoal agent and used in clinical practice for treatment of infectious gastroenteritis [24, 25], has emerged as a new broad-spectrum antiviral drug [26, 27]. We have previously reported that NTZ and its active circulating-metabolite tizoxanide (TIZ) are effective against different RNA viruses including influenza and parainfluenza viruses, hepatitis C and rotavirus infection *in-vitro* [28–31], as well as in clinical studies [32–34]. Interestingly nitazoxanide was also found to be effective against different animal and human coronaviruses, including MERS-CoV [26,35,36] and, more recently, SARS-CoV-2 *in-vitro* [37–39] and in COVID-19 patients [39–42].

While the molecular mechanism of its broad-spectrum antiviral activity has not been fully elucidated, NTZ was shown to inhibit the replication of influenza and parainfluenza viruses by a novel mechanism, impairing terminal glycosylation and intracellular trafficking of the class-I viral fusion glycoproteins influenza hemagglutinin and paramyxovirus F spike proteins [28, 29]. We therefore explored the possibility that NTZ may also affect SARS-CoV-2 spike maturation, which is critical for SARS-CoV-2 host recognition, penetration and pathogenesis [11]. Herein we report that nitazoxanide interferes with SARS-CoV-2 S-glycoprotein biogenesis, hampering its maturation at an endoglycosidase H (Endo-H)-sensitive stage, and hindering fusion and syncytium-forming ability of the original Wuhan spike, as wells as the Alpha, Beta, Gamma and Delta S-variants in human cells.

## RESULTS

### Expression and characterization of furin-dependent cleavage of SARS-CoV-2 S protein in human lung cells

Because of its fundamental role in virus infectivity and transmission, the SARS-CoV-2 spike protein structure and function have been extensively studied [7,10,43]; however, little is currently known about the SARS-CoV-2 S (SARS-2 S) biogenesis and maturation pathway.

In order to characterize SARS-CoV-2 S synthesis, oligomerization and furin-mediated cleavage as compared to SARS-CoV and MERS-CoV spike proteins in human lung cells, we have utilized two different expression plasmids, containing the full length SARS-CoV-2 spike gene ORF untagged (VG40589-UT, SARS-2 S) or harboring an antigenic tag (Flag) at the C-terminal (VG40589-CF, SARS-2 S-CF), as well as two plasmids containing the full length human SARS coronavirus (VG40150-CF) and MERS coronavirus (VG40069-CF) spike proteins both harboring a Flag-tag at the C-terminal.

We first asked whether SARS-2 S is robustly expressed in human lung alveolar type II-like epithelial A549 cells, as compared to human HEK293T cells, commonly used for experimentation because of their high transfectability; we also analyzed whether there is evidence for proteolytic processing of the S protein in lung cells, since it has been shown that SARS-2 S is cleaved by host cell proteases at the S1/S2 cleavage site in HEK293T cells [8].

Immunoblot analysis of A549 cells expressing the untagged SARS-2 S protein revealed a band with a molecular weight of 180 kDa expected for unprocessed S protein (S0) (Fig. 1C); a band with a size expected for the spike S1 subunit (∼110 kDa) [44] was also observed at all times tested. In addition to A549 cells, similar results were obtained in human normal lung MRC-5 fibroblasts (Fig. 1D). In A549 cells expressing the SARS-2 S protein with a C-terminal antigenic tag, in addition to the unprocessed S0, a band with a size expected for the S2 subunit (∼90 kDa) [45] was also observed when anti-Flag antibodies were utilized (Fig. 1E). These results suggest efficient proteolytic processing of SARS-2 S protein during biogenesis in human pulmonary cells, in keeping with the presence of several arginine residues at the S1/S2 cleavage site.

To confirm the involvement of furin in S protein processing during biogenesis, we analyzed whether the furin irreversible inhibitor decanoyl-RVKR-CMK (RVKR) blocks SARS-2 spike protein processing in our model. As shown in Fig. 1F,G,H, RVKR inhibited SARS-2 S processing in a concentration-dependent manner, confirming that the SARS-CoV-2 spike cleavage is mediated by furin; RVKR also inhibited the processing of MERS S protein, which is known to depend on furin [46], while it had no effect on SARS S protein expression (Fig. 1I), as expected.

It is interesting to note that inhibition of furin activity by RVKR did not prevent SARS-2 spike trimerization (Fig. 1H), indicating that S1/S2 cleavage is not required for oligomerization in lung cells.

### Nitazoxanide alters SARS-CoV-2 S protein biogenesis in human cells

We have previously shown that nitazoxanide (NTZ, Fig. 2A) inhibits the replication of influenza and parainfluenza viruses by impairing terminal glycosylation and intracellular trafficking of the class-I viral fusion glycoproteins influenza hemagglutinin and paramyxovirus fusion spike proteins [28, 29]. To investigate whether NTZ may also affect SARS-CoV-2 spike glycoprotein expression, human lung (A549 and MRC-5), colon (HCT116), liver (Huh7) and kidney (HEK293T) cells were transiently transfected with the SARS-2 S or SARS-2 S-CF plasmids and, after 4h, were treated with different concentrations of NTZ (Fig. 2B). As shown in Fig. 2C and Fig. S1A, NTZ did not inhibit S protein expression and did not interfere with the S1/S2 cleavage; however, treatment with NTZ at concentrations higher than 0.1 μg/ml caused an evident alteration in the molecular mass of both the S0 precursor and the S1/S2 subunits. This effect persisted for at least 24h (Fig. S1B) with no change in cell viability, as shown by MTT assay (NTZ LD_50_ >50 μg/ml). Interestingly NTZ treatment provoked a similar alteration of SARS-CoV and MERS-CoV spike glycoproteins exogenously expressed in A549 cells (Fig. 2D). Notably, the NTZ-mediated spike alteration was not affected by the presence of single and/or multiple mutations in the S protein, including the aspartic acid-to-glycine substitution at position 614 (D614G) (Fig. 2E), an S-protein globally-predominant mutation that was found to promote SARS-CoV-2 transmission in humans [47], and the N501Y, E484K and K417N mutations (Fig. S2A) that have been recently described in the B.1.1.7, P.1 and B.1.351 variants being monitored (VBMs) (https://www.cdc.gov/coronavirus/2019-ncov/variants/variant-info.html).

**Figure 2.**
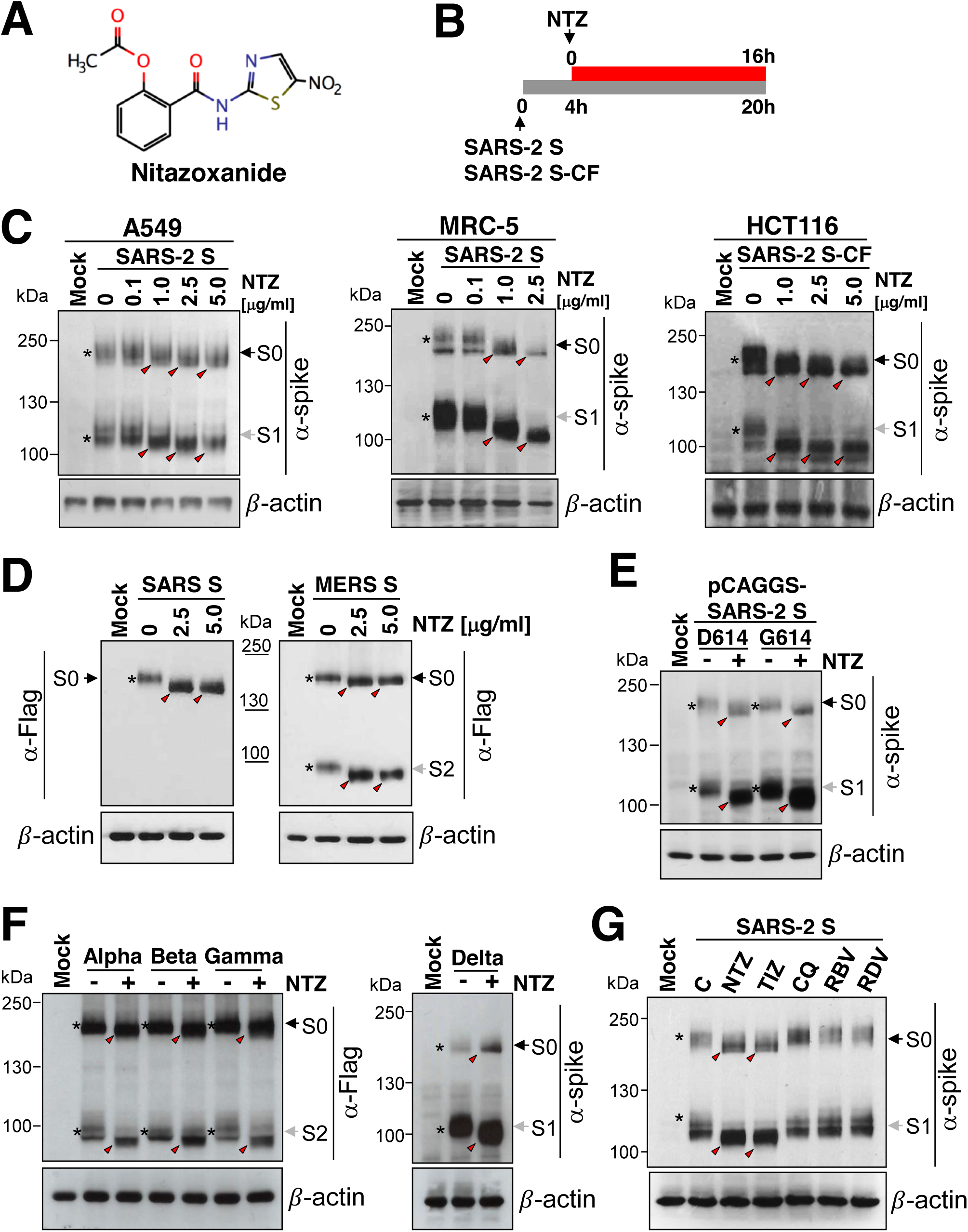
Nitazoxanide affects SARS-CoV-2 spike biogenesis in human cells. (**A**) Structure of nitazoxanide (NTZ). (**B**) Schematic representation of the experimental design. (**C**) Human lung (A549 and MRC-5) and colon (HCT116) cells were transiently transfected with the SARS-CoV-2 spike (SARS-2 S) (A549, MRC5) or C-terminal Flag-tagged SARS-CoV-2 spike (SARS-2 S-CF) (HCT116) constructs, or empty vector (Mock) and, after 4h, were treated with different concentrations of NTZ. After 16h, WCE were analyzed for levels of S protein by IB using anti-spike antibodies. (**D**) Levels of SARS-CoV and MERS-CoV S proteins were determined by IB using anti-Flag antibodies in WCE from A549 cells transiently transfected with C-terminal Flag-tagged SARS-CoV spike (SARS S) or MERS-CoV spike (MERS S) constructs or empty vector and treated with different concentrations of NTZ for 16h. (**E,F**) A549 cells were transiently transfected with C-terminal Flag-tagged pCAGGS-SARS2-S-D614 (D614) or pCAGGS-SARS2-S-G614 (G614) SARS-CoV-2 spike plasmids (**E**) or with C-terminal Flag-tagged pReceiver-M39 UK EX-CoV242-M39 (Alpha), EX-CoV244-M39-GS (Beta) and EX-CoV245-M39-GS (Gamma) or pCMV3-SARS-CoV-2-Spike (S1+S2)-long(B.1.617.2) (Delta) plasmids (**F**) or empty vector, and treated with NTZ (2.5 µg/ml) or vehicle (-). After 16h, WCE were analyzed for levels of S protein by IB using anti-spike (**E,F**) or anti-Flag (**F**) antibodies. (**G**) Immunoblot of S protein (α-spike) levels in A549 cells transfected with the SARS-2 S construct for 4h and treated with 2.5 μg/ml NTZ, 2.5 μg/ml tizoxanide (TIZ), 10 μM chloroquine (CQ), 7.5 μM ribavirin (RBV), 7.5 μM remdesivir (RDV) or vehicle for 16h. (**C-G**) Black arrows indicate bands corresponding to uncleaved S proteins (S0), whereas gray arrows indicate bands corresponding to the S1 or S2 subunits. The slower- and faster-migrating forms of the uncleaved S protein or cleaved subunits are identified by asterisks and red arrowheads respectively.

In addition, similar results were obtained using plasmids encoding the Alpha B.1.1.7 (UK), Beta B.1.351 (South Africa) and Gamma P.1 (Brazil) VBMs, and the Delta B.1.617.2 variant of concern (VOC) spike-glycoproteins exogenously expressed in A549 cells (Fig. 2F). Finally, it should be noted that, in addition to human cells, nitazoxanide was found to affect SARS-CoV-2 spike biogenesis also in monkey (Vero E6), rodent (murine L929 fibroblasts) and bat lung epithelial cells Tb-1 Lu cells (Fig. S1C-E), indicating a general effect on mammalian cells.

The NTZ-induced alteration in the S glycoprotein electrophoretic mobility pattern was mimicked by the NTZ active metabolite, tizoxanide (Fig. S3A), but not by other antiviral drugs, including chloroquine (10 μM), ribavirin (7.5 μM) and remdesivir (7.5 μM) (Fig. 2G).

Whereas the antiviral activity of nitazoxanide against SARS-CoV-2 replication has been reported *in-vitro* and in clinical settings [37–42], the effect of tizoxanide on SARS-CoV-2 infection is not well established. Therefore the antiviral activity of tizoxanide was examined in a model of SARS-CoV-2 (SARS-CoV-2/England/2/2020) infection in Vero E6 cells. The results, shown in Fig. S3B, demonstrate that the active metabolite of nitazoxanide is effective in inhibiting SARS-CoV-2 replication *in-vitro* with an EC_50_ of 2.9 μM at 24h post infection.

### Nitazoxanide does not affect SARS-CoV-2 spike stability and oligomerization, but impairs its maturation at an Endo-H sensitive stage

To investigate whether NTZ treatment affects SARS-CoV-2 S protein stability, A549 cells were transfected with the SARS-2 S construct and treated with NTZ (2.5 μg/ml) for 16h, at which time protein synthesis was arrested by adding cycloheximide (100 μg/ml). At different times after cycloheximide treatment S protein levels were determined in whole-cell extracts by Western blot and quantitated by scanning densitometry. The results, shown in Fig. 3A, indicate that S protein degradation rate (with a half-life of 7.5h) is not significantly affected by the thiazolide. Comparable levels of S-protein were also detected by confocal microscopy in A549 cells untreated or treated with nitazoxanide (5 μg/ml) for 16h (Fig. 3B). Notably, nitazoxanide did not prevent the formation of SARS-2 spike dimers and trimers, which, however, were characterized by an evident alteration in electrophoretic mobility, reflecting the monomer modification (Fig. 3C).

**Figure 3.**
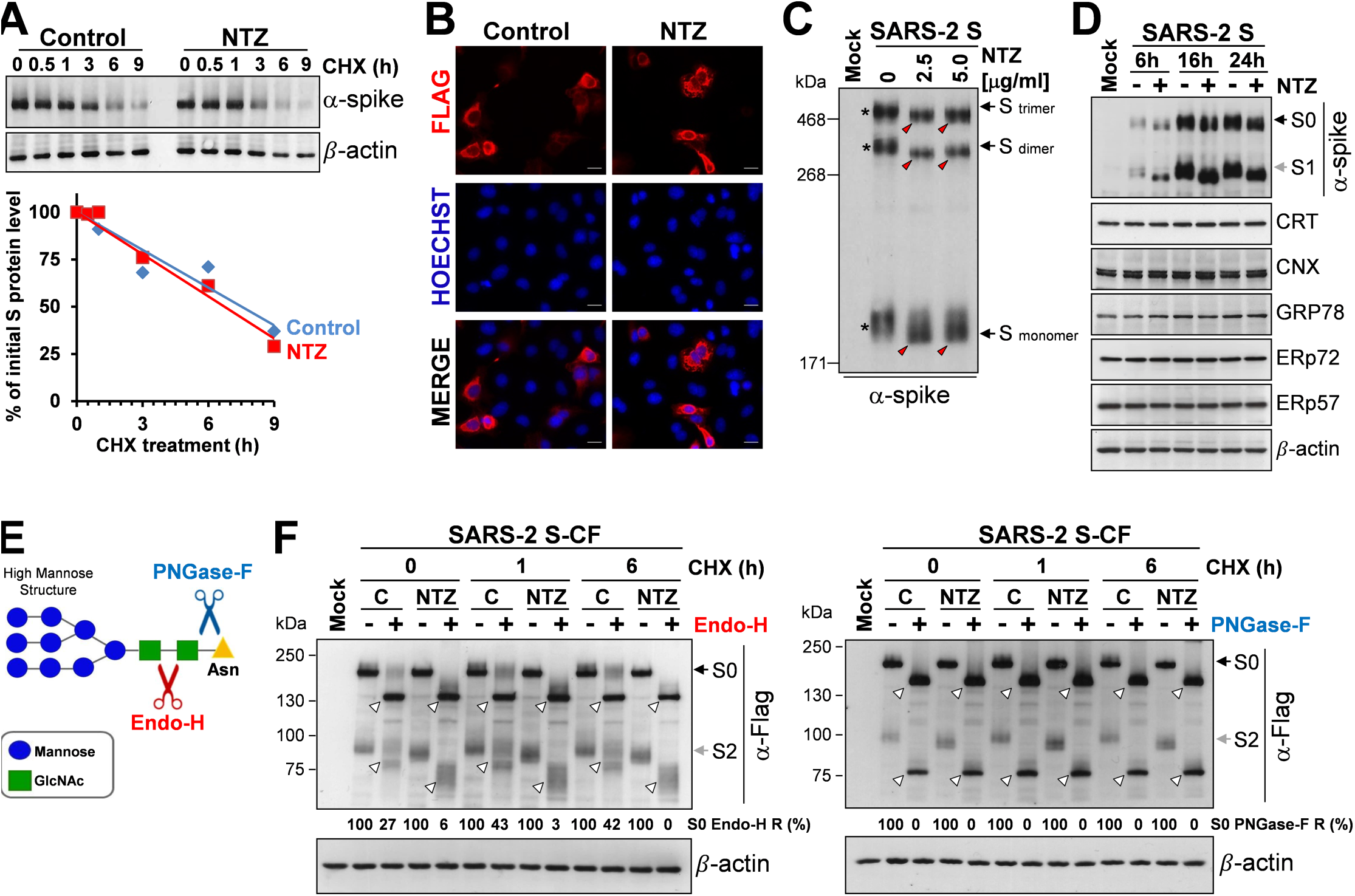
Nitazoxanide does not affect SARS-CoV-2 spike stability and oligomerization, but impairs its maturation at an Endo-H sensitive stage. (**A**) A549 cells were transfected with the SARS-CoV-2 spike (SARS-2 S) construct for 4h and treated with NTZ (2.5 μg/ml) or vehicle. After 16h, cycloheximide (CHX, 100 μg/mL) was added (time 0), and WCE were analyzed for S protein levels at different times after CHX addition by IB using anti-spike antibodies (top) and quantitated by scanning densitometry. Relative amounts of S protein were determined after normalizing to β-actin (bottom). The plot of percentage of initial S protein levels in control (blue) and NTZ-treated (red) cells versus time is shown. (**B**) Confocal images of Flag-tagged S protein (red) in A549 cells transfected with the C-terminal Flag-tagged SARS-CoV-2 spike (SARS-2 S-CF) construct for 4h and treated with NTZ (5 μg/ml) or vehicle for 16h. Nuclei are stained with Hoechst (blue). Merge images are shown. Scale bar, 20 µm. Data from a representative experiment of three with similar results are shown. (**C**) Gel electrophoresis (4% polyacrylamide) of WCE from A549 cells transfected with SARS-2 S construct or empty vector and treated with different concentrations of NTZ for 16h. The different forms (trimers, dimers and monomers) of the S protein were visualized by IB with anti-spike antibodies. The slower- and faster-migrating forms of S protein are identified by asterisks and red arrowheads respectively. (**D**) IB of S protein (α-spike), calreticulin (CRT), calnexin (CNX), GRP78 (anti-KDEL), ERp72 and ERp57 in WCE from A549 cells transfected with the SARS-2 S construct or empty vector for 4h and treated with 2.5 μg/ml NTZ (+) or vehicle (-) for different times. (**E**) Diagram of substrate specificities of endoglycosidase H (Endo-H) and peptide-N-glycosidase F (PNGase-F). Red and blue scissors represent the cleavage sites of Endo-H (red) and PNGase F (blue), respectively. Mannose (blue circles), N-acetylglucosamine (GlcNAc, green squares) and asparagine (Asn, yellow triangle) residues are shown. (**F**) A549 cells transfected with C-Flag tagged SARS-CoV-2 spike (SARS-2 S-CF) construct or empty vector (Mock) for 4h were treated with NTZ (2.5 µg/ml) or vehicle. After 16h, CHX (100 µg/mL) was added (time 0); at different times after CHX addition proteins were digested with Endo-H or PNGase-F (+) or left untreated (-) and processed for IB analysis using anti-Flag antibodies and quantitated by scanning densitometry. The Endo-H- or PNGaseF-cleaved faster-migrating S and S2 forms are indicated by white arrowheads. The percentage of Endo-H-resistant (Endo-H R) S protein in the different samples is indicated. Black arrows indicate bands corresponding to uncleaved S protein (S0), whereas gray arrows indicate bands corresponding to the S1 or S2 subunits.

Although it is known that glycosylation plays an essential role in establishing coronavirus spike proteins bioactive conformation, stability, tropism and fusion activity [10,13,14], presently there is little information on the effect of glycosylation inhibitors on SARS-CoV-2 spike biogenesis. The thiazolide-induced SARS-2 spike alteration was then compared to the effect of the *N-*glycosylation inhibitor tunicamycin. Interestingly, non-glycosylated spike protein was not detected in the soluble fraction of tunicamycin-treated cells (Fig. S4A, top), an effect that could not be rescued by treatment with the proteasome inhibitor bortezomib [48], suggesting that non-glycosylated proteins are not promptly degraded via ERAD (ER-associated degradation), a sophisticated process for ER-to-cytosol dislocation and ubiquitin/proteasome-mediated degradation of target-proteins [49]. In fact, immunomicroscopy studies revealed the formation of large S protein aggregates in tunicamycin-treated cells (Fig. S4B); in addition, the presence of a low-molecular weight band, corresponding to the non-glycosylated spike protein (∼130 kDa), was detected in the insoluble fractions of tunicamycin-treated cell lysates (Fig. S4A, bottom). Similar results were previously reported for the spike protein of transmissible gastroenteritis virus (TGEV) CoV [50].

Glycosylation of coronavirus S proteins, like other cell surface glycoproteins, is initiated in the endoplasmic reticulum, adding the “high mannose” oligosaccharides; the mannose-rich sugar component is processed in the Golgi apparatus, and terminal glycosylation occurs in the *trans* cisternae of the Golgi apparatus [51].

We have previously shown that nitazoxanide blocks the maturation of the influenza virus hemagglutinin at a stage preceding resistance to digestion by endoglycosidase H (Endo-H), an enzyme that removes *N*-linked carbohydrate chains that have not been terminally glycosylated (Fig. 3E) [52], thus impairing HA intracellular trafficking [28]. To obtain insights on the effect of NTZ on SARS-2 S maturation, we therefore investigated whether nitazoxanide could affect S protein terminal glycosylation. SARS-CoV-2 S protein aliquots from NTZ-treated or control cells were subjected to digestion with Endo-H or with PNGase-F, an enzyme that removes all *N*-glycans [53], up to 6 h after cycloheximide treatment and then analyzed by immunoblot. As expected, S proteins from both untreated and NTZ-treated cells were sensitive to PNGase-F digestion (Fig. 3F). On the other hand, a fraction (approximately 42-43% between 1 and 6h after cycloheximide treatment) of the SARS-CoV-2 S protein was found to be terminally glycosylated becoming Endo-H-resistant in control cells under the conditions described; notably, S protein from NTZ-treated cells remained instead sensitive to digestion with the glycosidase up to 22h after synthesis (Fig. 3F). Because acquisition of Endo-H resistance is a marker for transport into the *cis* and *middle* Golgi compartments [52], these results indicate that nitazoxanide, as in the case of influenza HA [28], may impair S protein trafficking between the ER and the Golgi complex. Of note, treatment with nitazoxanide did not significantly alter the expression of calnexin or other ER proteins, including calreticulin, ERp57, ERp72 and the ER-stress regulated protein GRP78, suggesting that the drug is not triggering the host ER-stress response (Fig. 3D). It should also be noted that, at the effective concentration, nitazoxanide does not alter the expression of the other SARS-CoV-2 structural proteins N, M and E when expressed alone (Fig. S5A) or in combination (Fig. S5B). It is important to note that the presence of large amounts of the nucleocapsid, envelope and in particular the membrane protein, which is known to interact with the S glycoprotein [54], did not prevent the NTZ-induced spike alteration (Fig. S5B).

Finally, the thiazolide-induced SARS-2 spike alteration was also compared to the effect of different glycoprotein maturation inhibitors, including the α-glucosidases inhibitor castanospermine, the ER-Golgi protein trafficking inhibitor brefeldin A, and the mannosidase-I inhibitors deoxymannojirimicin (DMJ) and kifunensine, which was recently shown to alter SARS-CoV-2 spike processing and infectivity [55]. As shown in Fig. S4C, neither α-glucosidases nor mannosidase inhibitors caused an alteration in molecular mass comparable to nitazoxanide, suggesting a different activity of the drug. Interestingly, brefeldin A treatment, differently from nitazoxanide, inhibited S protein cleavage (Fig. S4C).

These results indicate that, as in the case of influenza HA and parainfluenza fusion proteins, nitazoxanide treatment results in hampering SARS-CoV-2 spike maturation.

### Nitazoxanide treatment prevents SARS-CoV-2 S pseudovirions infection

Since biochemical assays using purified viral components do not provide insights on the S protein function as it occurs on the virion membrane, we engineered pseudotyped retroviral (Murine Leukemia Virus, MLV) as well as third-generation lentiviral (pLV) particles to express the full-length SARS-CoV-2 S protein together with a luciferase (LUC) or green fluorescent protein (GFP) reporter to monitor infection (Fig. 4A).

**Figure 4.**
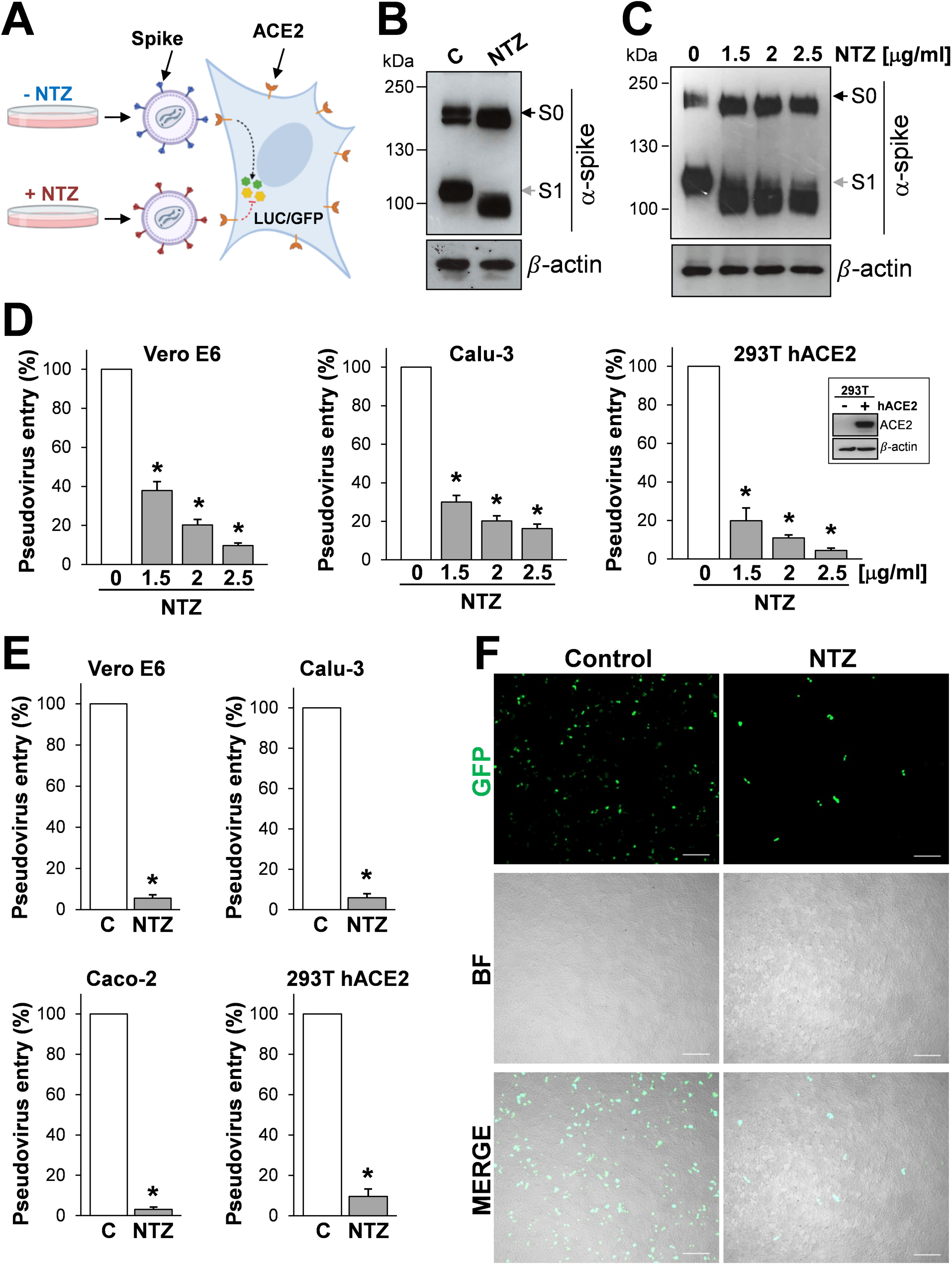
Nitazoxanide treatment inhibits SARS-CoV-2 S pseudovirus infectivity. (**A**) Diagram of SARS-CoV-2 pseudovirus luciferase (LUC) and green fluorescent protein (GFP) assay. SARS-CoV-2 spike pseudotyped lentiviral (LV) or retroviral (MLV) particles carrying a reporter gene are produced in the presence of NTZ or vehicle (C) and used to infect different types of human cells, which will express LUC or GFP upon infection. (**B,C**) HEK293T cells transfected with plasmids for production of SARS-CoV-2 spike pseudotyped-LUC LV particles (**B**) or SARS-CoV-2 spike pseudotyped-LUC MLV particles (**C**) (see ’Materials and Methods’), were treated with 2.5 μg/ml (**B**) or different concentrations (**C**) of NTZ. At 48h post-transfection, pseudovirus-containing supernatants were collected for infection of different types of cells (see panels **D,E**), whereas SARS-CoV-2 S expression was detected in WCE of HEK293T pseudovirus-producing cells by IB using anti-spike antibodies. (**D,E)** Vero E6, Calu-3, Caco-2 and hACE2-expressing HEK293T (293T hACE2) cells were infected with SARS-CoV-2 spike-pseudotyped MLV (**D**) or LV (**E**) particles obtained as described in **B,C**, and pseudovirus entry was analyzed by measuring LUC activities 72h post-infection. hACE2 protein levels in HEK293T WT (-) and hACE2 (+) cells are shown in **D**. Data are expressed as percent of untreated control. Error bars indicate means±SD. *P<0.05; ANOVA (**D**), Student’s *t-*test (**E**). **(F)** SARS-CoV-2 spike-pseudotyped GFP lentiviral particles produced as described in **B** were used to infect 293T hACE2 cells. Pseudovirus-infected cells were visualized by fluorescence microscopy. Bright field (BF) and merge images are shown. Scale bar, 200 µm.

SARS-CoV-2 S-pseudotyped particles were generated in producer HEK293T cells treated with different concentrations of nitazoxanide or control diluent for 48h and then used to infect different types of cells, including monkey Vero E6 cells, that are commonly used in the study of SARS-CoV-2 entry due to their high susceptibility to infection, as well as human airway epithelial Calu-3 cells and intestinal epithelial Caco-2 cells, which both express the ACE2 receptor on their apical surface [56]; in addition, as HEK293T cells were not susceptible to infection by SARS-CoV-2 S-pseudotyped viruses [45], we generated HEK293T cells stably expressing the human ACE2 receptor (293T hACE2 cells) for these studies.

As shown in Fig. 4, nitazoxanide treatment caused the expected SARS-CoV-2 spike alteration in HEK293T producer cells (Fig. 4B,C) and resulted in a dramatic decrease of both MLV (Fig. 4D) and pLV (Fig. 4E) SARS-CoV-2 S-pseudotyped particles infectivity in all cell lines examined, as determined quantitatively by luciferase levels detection. Similar results were obtained when HEK293T producer cells were treated with the NTZ metabolite tizoxanide (Fig. S3C,D).

A marked decrease in infectivity of pseudovirions containing NTZ-modified proteins was also assessed qualitatively by fluorescence microscopy in 293T hACE2 cells infected with GFP-expressing SARS-CoV-2 S-pseudotyped LV particles (Fig. 4F). Similar results were obtained when GFP-expressing SARS-CoV-2 S-pseudotyped LV particles were used to infect Caco-2 hACE2 cells, intestinal epithelial cells engineered to express high levels of the human ACE2 receptor (Fig. S6).

### Nitazoxanide inhibits SARS-CoV-2 S-mediated cell-cell fusion and syncytia formation

Besides fusion mediated by virions, SARS-CoV-2 S proteins present at the plasma membrane, contrary to the S protein of the related coronavirus SARS-CoV [57], can trigger receptor-dependent formation of syncytia, which have been observed not only in cell cultures but also in tissues from individuals infected with SARS-CoV-2 [15, 58].

To address the question whether the alteration in SARS-CoV-2 S glycoprotein maturation observed in nitazoxanide treated cells may affect cell-cell fusion and syncytia formation, we utilized two different approaches.

In the first approach monkey Vero E6 cells and human Huh7 cells, both naturally expressing ACE2 receptors on the membrane surface, transiently transfected with the Flag-tagged SARS-CoV-2 or SARS-CoV (Huh7) spike plasmids were treated with nitazoxanide or vehicle, and processed for immunofluorescence microscopy using an anti-Flag antibody to visualize syncytia formation after 36h. Consistently with previous reports [57], expression of the SARS-CoV S-glycoprotein did not cause syncytia formation in the absence of trypsin in ACE2-expressing Huh7 cells, whereas SARS-CoV-2 spike expression resulted in the formation of large multi-nucleated syncytia in the same model (Fig. 5A). Interestingly, treatment with nitazoxanide, whereas it did not inhibit SARS-CoV-2 spike expression, markedly decreased the number and size of syncytia in both Vero E6 cells (Fig. 5B) and Huh7 cells (Fig. S7), indicating that nitazoxanide may prevent SARS-CoV-2 S-mediated fusion between adjacent cells (fusion ’from within’).

**Figure 5.**
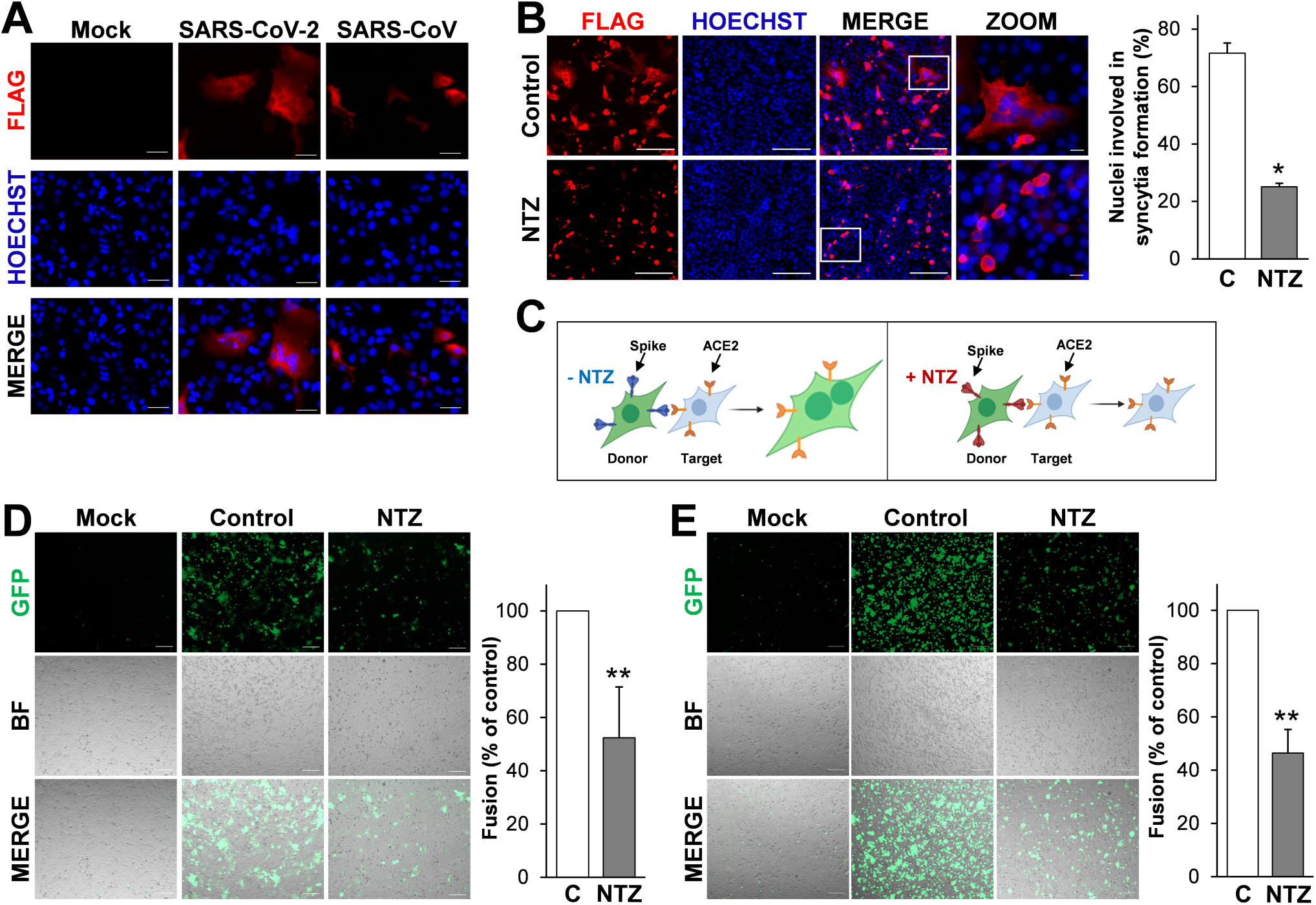
Nitazoxanide inhibits SARS-CoV-2 S-mediated cell-cell fusion. (**A**) Syncytia formation by Huh7 cells transfected with the C-terminal Flag-tagged SARS-CoV-2 or SARS-CoV spike constructs or empty vector (Mock) for 40h were analyzed by immunofluorescence microscopy using anti-Flag antibodies (red). Nuclei are stained with Hoechst (blue). Merge images are shown. Scale bar, 20 µm. (**B**Vero E6 cells transfected with the SARS-2 S-CF construct for 4h and then treated with NTZ (5 μg/ml) or vehicle (Control) for 36h were analyzed by immunofluorescence microscopy using anti-Flag antibodies (red). Nuclei are stained with Hoechst (blue). Merge and zoom images are shown. Scale bar, 200 µm (zoom, 20 µm). The number of nuclei involved in syncytia formation is expressed as percent of total nuclei in transfected cells in the same sample (right panel). Data represent the mean±SD of 5 fields from three biological replicates. *P<0.05; Student’s *t*-test. (**C)** Schematic representation of the donor-target cell-cell fusion assay. (**D,E)** HEK293T cells co-transfected with plasmids encoding SARS-CoV-2 S and GFP for 4h and treated with NTZ (5 μg/ml) or vehicle (Control) for 36h were overlaid on Vero E6 (**D**) or Huh7 (**E**) cell monolayers. After 4h, cell-cell fusion was assessed by fluorescence microscopy. Bright field (BF) and merge images are shown. Scale bar, 200 µm. Cell-cell fusion was determined and expressed as percentage relative to control (right panels). Data represent means±SD of three replicates. **P<0.01; Student’s *t-*test.

In a second type of approach, we investigated whether the drug may prevent cell-cell fusion between different cell types. HEK293T cells were co-transfected with plasmids encoding the Flag-tagged SARS-CoV-2 S glycoprotein and GFP and, after 4h, treated with nitazoxanide or vehicle (Fig. 5C). At 40h post transfection cells were detached and overlaid on an 80% confluent monolayer of target Vero E6 and Huh7 cells at a ratio of approximately one S-expressing cell to three receptor-expressing cells. After 4h, co-cultures were processed for fluorescence microscopy to visualize syncytia formation, and the extent of cell–cell fusion was determined as described [59]. As shown in Fig. 5D,E, also in this case treatment with nitazoxanide markedly decreased the number and size of syncytia in both Vero E6 cells (Fig. 5D) and Huh7 cells (Fig. 5E), confirming that nitazoxanide may prevent SARS-CoV-2 S-mediated fusion between different types of cells.

Formation of syncytia by pneumocytes expressing SARS-CoV-2 RNA and S proteins was observed post-mortem in lung tissues of COVID-19-infected patients [58]. We therefore investigated whether nitazoxanide treatment may prevent SARS-CoV-2 S-mediated fusion in human lung alveolar type II-like A549 cells. As these cells are known to possess low levels of ACE2 [8], we generated A549 cells stably expressing the human ACE2 receptor (A549 hACE2 cells) for these studies. As shown in Fig. 6A, exogenous ACE2 expression resulted in a dramatic increase in SARS-CoV-2 S-mediated fusion between adjacent lung cells, confirming the important role of ACE2 in virus-directed syncytia formation. Consistently with the results obtained in other cell types, treatment with nitazoxanide markedly decreased the number and size of syncytia in A549 hACE2 cells (Fig. 6A). Similar results were obtained when A549 hACE2 cells were used as target cells in the cell-cell fusion co-culture model, using HEK293T cells co-transfected with plasmids encoding the Flag-tagged SARS-CoV-2 S-glycoprotein and a GFP reporter as donor cells (Fig. 6B). NTZ-induced inhibition of cell-cell fusion was dose-dependent, with a >90% suppression of syncytia formation at the concentration of 10 μg/ml (Fig. S8A).

**Figure 6.**
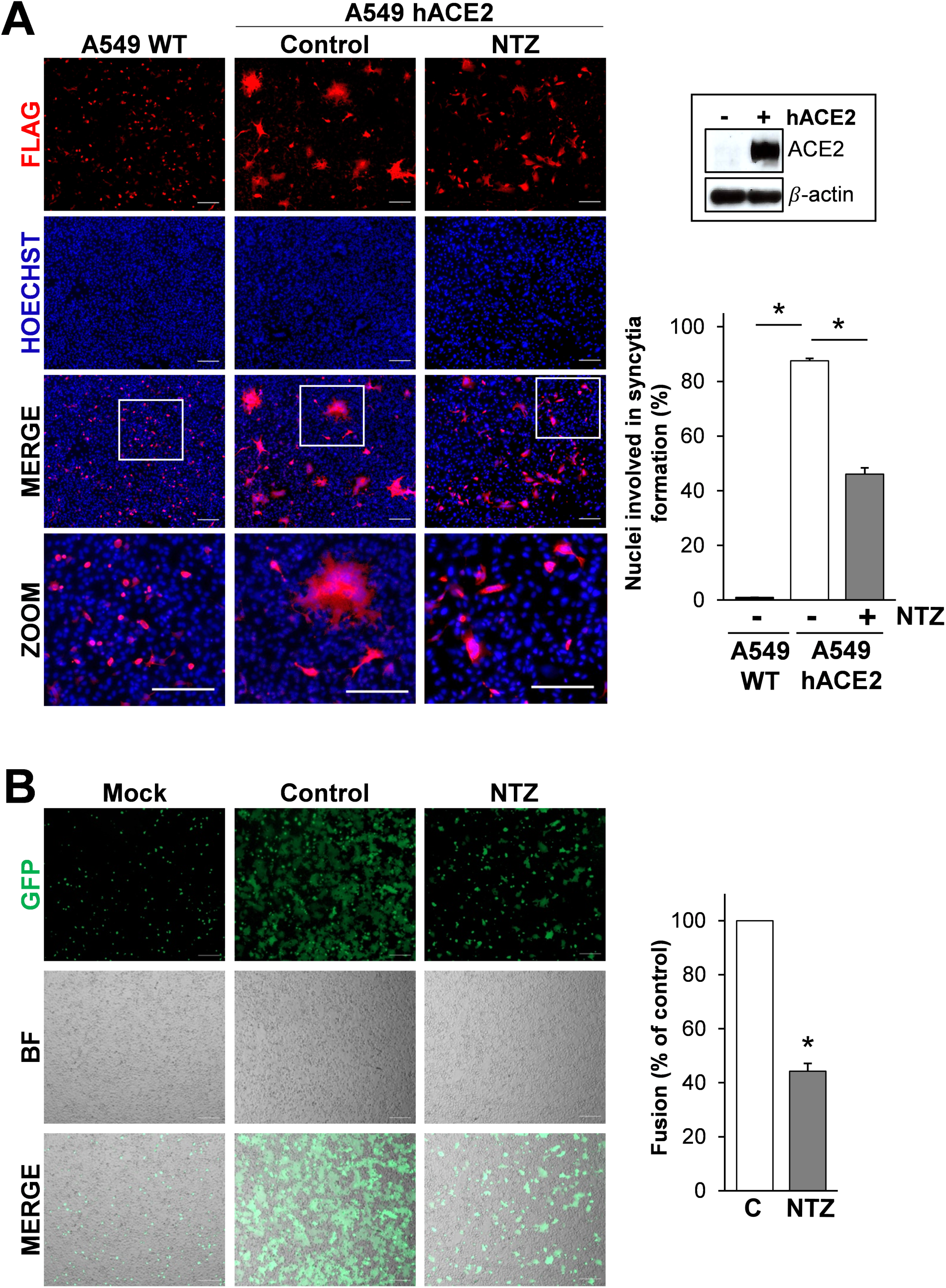
Nitazoxanide treatment hinders SARS-CoV-2 S-mediated cell fusion in human lung epithelial A549 cells. (**A**) A549 cells stably expressing human ACE2 (A549 hACE2) or wild-type (WT) were transiently transfected with the SARS-2 S-CF construct for 4h and treated with NTZ (5 μg/ml) or vehicle (Control) for 36h. Immunofluorescence analysis was performed using an anti-Flag antibody (red). Nuclei are stained with Hoechst (blue). Merge and zoom images are shown. Scale bar, 200 µm (zoom, 200 µm). hACE2 protein levels in A549 WT (-) and A549 hACE2 (+) cells are shown. The number of nuclei involved in syncytia formation is expressed as percent of total nuclei in transfected cells in the same sample (right panel). Data represent the means±SD of 5 fields from three biological replicates. *P<0.05; ANOVA. (**B)** HEK293T cells co-transfected with plasmids encoding SARS-CoV-2 S and GFP for 4h and treated with NTZ (5 μg/ml) or vehicle (Control) for 36h were overlaid on A549 hACE2 cell monolayers. After 4h, cell-cell fusion was assessed by fluorescence microscopy (left panel). Bright field (BF) and merge images are shown. Scale bar, 200 µm. Cell-cell fusion was determined and expressed as percentage relative to control (right panel). Error bars indicate means±SD. *P<0.05; Student’s *t*-test.

It is known that the SARS-CoV-2 E and M proteins regulate processing and intracellular trafficking of the S glycoprotein, interfering with syncytia formation [54,60,61]; we therefore investigated whether co-expression of the structural proteins N, M and E with the spike glycoprotein would affect the ability of the drug to inhibit S-mediated cell-cell fusion. As shown in Fig. S5C, the presence of the N, M and E proteins did not prevent cell-cell fusion and did not hinder the ability of nitazoxanide to suppress syncytia formation in lung cells.

These results confirm that SARS-CoV-2 S-protein possesses high fusogenic activity and is able to trigger large syncytia formation in human lung cells, and indicate that the alteration in SARS-CoV-2 S glycoprotein maturation observed in nitazoxanide-treated cells impairs spike-driven cell-cell fusion and syncytia formation.

### Nitazoxanide inhibits the fusogenic activity of SARS-CoV-2 spike variants

Although coronaviruses are thought to mutate at a slower rate than most RNA viruses due to the presence of a unique viral proofreading machinery [1, 62], CoVs can also undergo recombination events [63]. A meta-analysis of genomes of SARS-CoV-2 strains found several mutations within the virion spikes circulating in a considerable percentage of the analyzed populations [21–23]. The most common mutation, now found in most of the global population, is an aspartic acid-to-glycine substitution at position 614 (D614G) in the S1 subunit of the spike protein [47]. The D614G S mutation has been shown to promote SARS-CoV-2 transmission in humans and to enhance functional S-protein incorporation into SARS-CoV-2 virus-like particles and pseudovirus particles, increasing pseudovirions infectivity [64].

To investigate whether nitazoxanide treatment was equally effective in inhibiting the infectivity of the SARS-CoV-2 D614G spike variant, we utilized the pCAGGS-SARS2-S-D614 S (wild type, SARS-2 D614) plasmid and the pCAGGS-SARS2-S-G614 S plasmid carrying the D614G mutation (SARS-2 G614), both containing a C-terminal antigenic Flag-tag, to engineer pseudotyped lentiviral particles expressing the full-length SARS-CoV-2 S protein together with a LUC reporter. SARS-CoV-2 D614G S-pseudotyped particles were generated in producer HEK293T cells treated with nitazoxanide (2.5 μg/ml) for 48h and then used to infect 293T hACE2 cells and intestinal epithelial Caco-2 hACE2 cells.

As shown in Fig. 7A, nitazoxanide treatment caused the expected SARS-CoV-2 D614G spike alteration in HEK293T producer cells and resulted in a dramatic decrease of SARS-CoV-2 D614G S pseudotyped particles infectivity in both 293T hACE2 and Caco-2 hACE2 cells (Fig. 7B).

**Figure 7.**
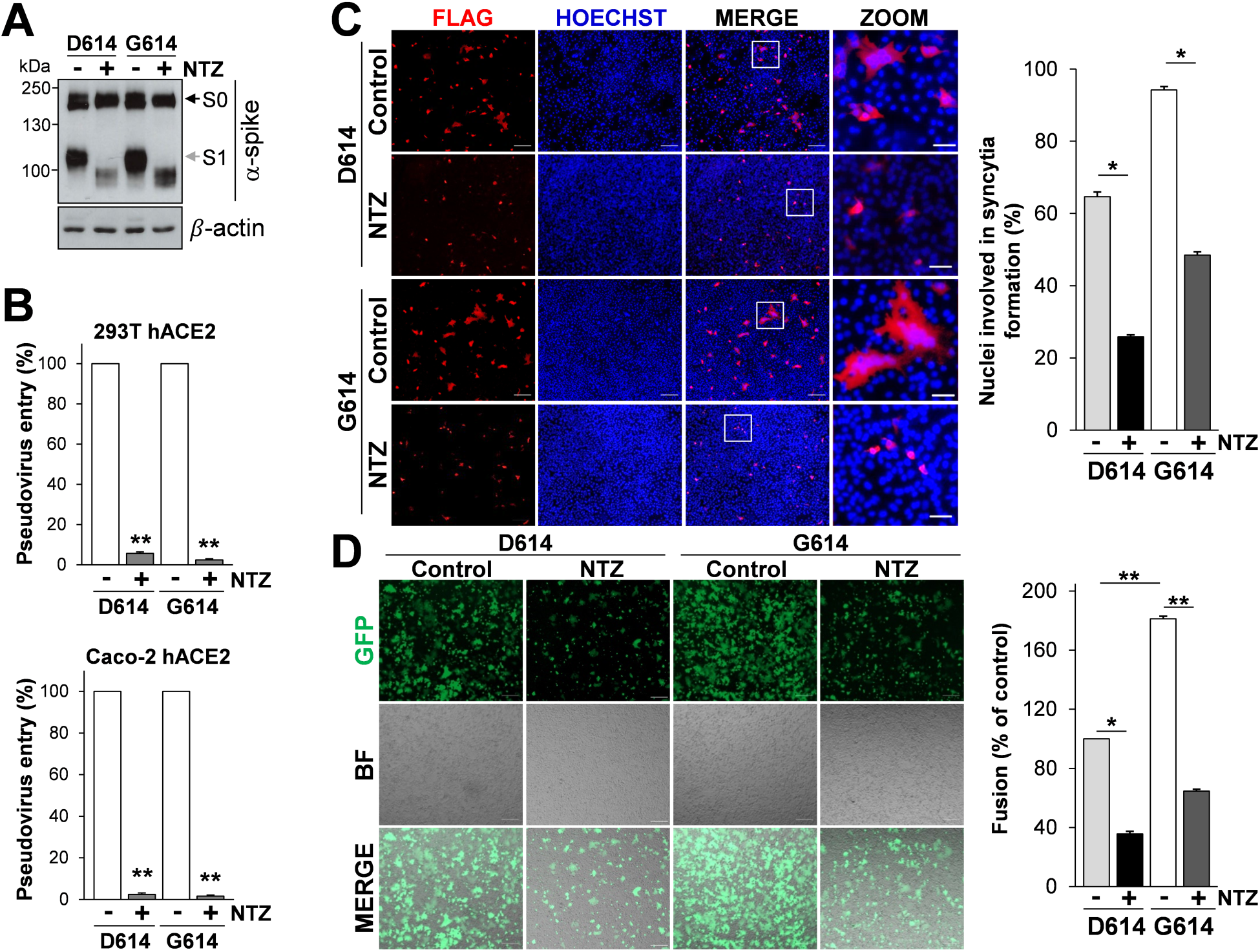
Nitazoxanide inhibits the fusogenic activity of the SARS-CoV-2 D614G spike variant. (**A,B**) HEK293T cells transfected with plasmids for production of SARS-CoV-2 D614 or G614 spike pseudotyped-LUC LV particles, as described in Fig. 4A, were treated with 2.5 μg/ml of NTZ (+) or vehicle (-). At 48h post-transfection, pseudovirus-containing supernatants were collected for infection of 293T hACE2 or Caco-2 hACE2 cells. SARS-CoV-2 S expression was detected in WCE of HEK293T pseudovirus-producing cells by IB using anti-spike antibodies (**A**). 293T hACE2 or Caco-2 hACE2 cells were infected with SARS-CoV-2 spike-pseudotyped LV-particles obtained as described above and pseudovirus entry was analyzed by measuring LUC activities 72h post-infection (**B**). Data are expressed as percent of relative untreated controls. Error bars indicate means±SD. **P<0.01; Student’s *t*-test. **(C)** A549 hACE2 cells were transiently transfected with the C-Flag tagged SARS-CoV-2 D614 or G614 spike constructs for 4h and treated with NTZ (5 μg/ml) or vehicle (Control) for 36h. Immunofluorescence analysis was performed using an anti-Flag antibody (red). Nuclei are stained with Hoechst (blue). Merge and zoom images are shown. Scale bar, 200 µm (zoom, 50 µm). The number of nuclei involved in syncytia formation is expressed as percent of total nuclei in transfected cells in the same sample (right panel). (**D)** HEK293T cells co-transfected with plasmids encoding the C-Flag tagged SARS-CoV-2 D614 or G614 spike and GFP for 4h, and treated with NTZ (5 μg/ml) or vehicle (Control) for 36h were overlaid on A549 hACE2 cell monolayers. After 4h, cell-cell fusion was assessed by fluorescence microscopy (left panel). Bright field (BF) and merge images are shown. Scale bar, 200 µm. Cell-cell fusion was measured and expressed as percentage relative to the D614 control (right panel). (**C,D**) Data represent means±SD of 5 fields from three replicates. *P<0.05, **P<0.01; ANOVA.

Next, human lung A549 hACE2 cells were transfected with the Flag-tagged SARS-CoV-2 D614 or G614 spike plasmids, treated with nitazoxanide or vehicle, and processed for immunofIuorescence microscopy using an anti-Flag antibody to visualize syncytia formation after 36h. As shown in Fig. 7C, expression of the SARS-CoV-2 G6k14 S glycoprotein was markedly more effective in causing the formation of large multi-nucleated syncytia in lung cells as compared to the wild-type protein. Treatment with nitazoxanide was found to significantly decrease the number and size of syncytia caused by the SARS-CoV-2 G614 spike variant (Fig. 7C).

Similar results were obtained when A549 hACE2 cells were used as target cells in the fusion co-culture model described above, using HEK293T cells co-transfected with plasmids encoding the SARS-CoV-2 S D614 or G614 spike variant together with a GFP reporter as donor cells (Fig. 7D). In addition to the G614 spike variant, nitazoxanide was found to significantly decrease the number and size of syncytia caused by SARS-CoV-2 spike proteins carrying single or multiple mutations including the N501Y, E484K and K417N substitutions (Fig. S2B,C) that, as indicated above, have been recently described in the B.1.1.7, P.1 and B.1.351 VBMs.

In the last year the variants that have major global health impact include Alpha (B.1.1.7), Beta (B.1.351), Gamma (P.1) VBMs and Delta (B.1.617.2) VOC variants. We therefore investigated whether nitazoxanide was effective in inhibiting the infectivity of these SARS-CoV-2 S variants. We utilized the EX-CoV242-M39 (Alpha), EX-CoV244-M39-GS (Beta), EX-CoV245-M39-GS (Gamma) and pCMV3-SARS-CoV-2-Spike (S1+S2)-long(B.1.617.2) (Delta) plasmids to engineer pseudotyped lentiviral particles expressing the full-length SARS-CoV-2 S proteins together with a LUC reporter, as reported above. SARS-CoV-2 Alpha, Beta, Gamma and Delta S-pseudotyped particles were generated in producer HEK293T cells treated with nitazoxanide (2.5 μg/ml) for 48h and then used to infect Caco-2 hACE2 cells. As shown in Fig. 8, nitazoxanide treatment resulted in a dramatic decrease in the infectivity of all SARS-CoV-2 S pseudotyped particles (Fig. 8A-D, I).

**Figure 8.**
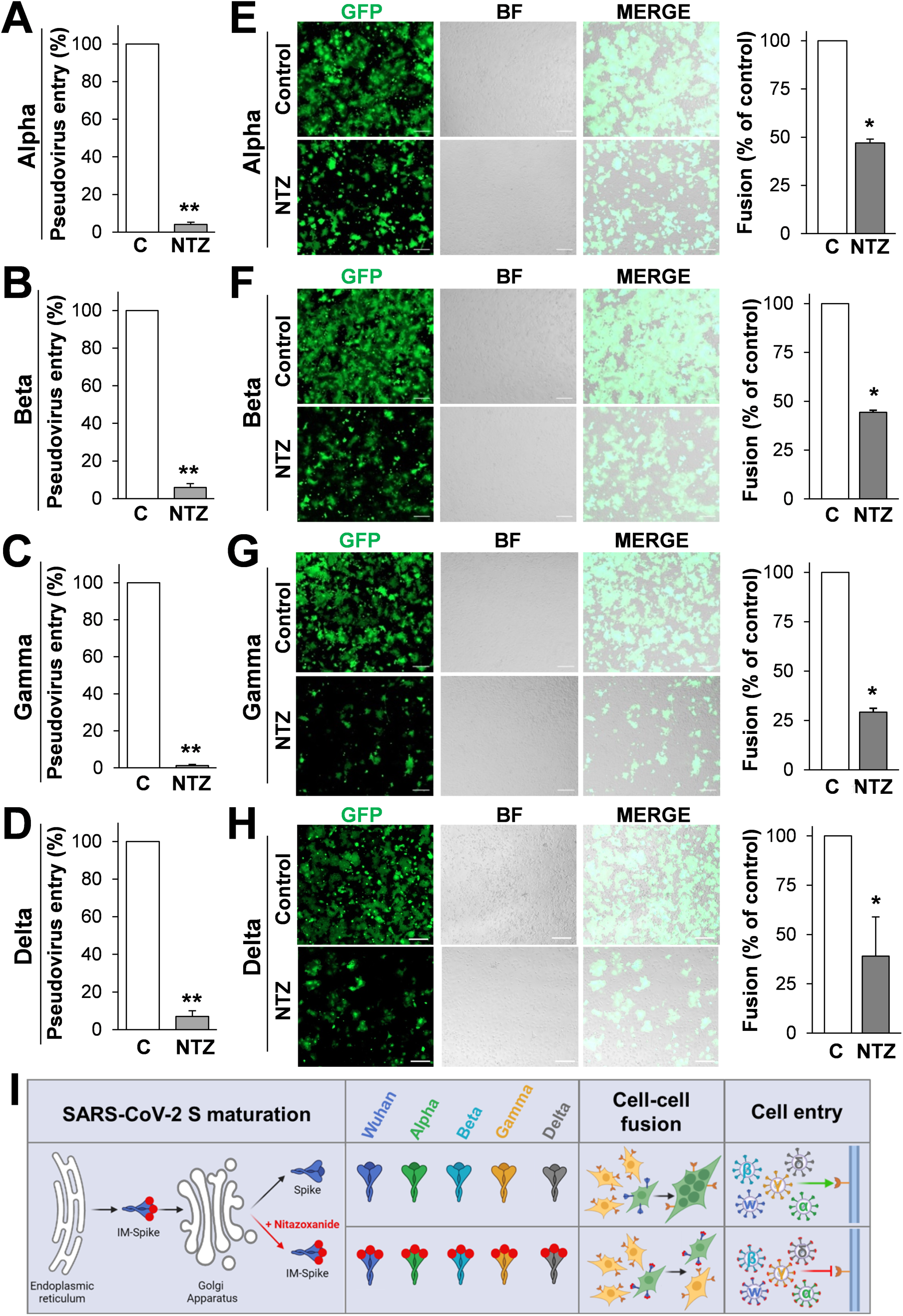
SARS-CoV-2 Alpha, Beta, Gamma and Delta spike variants pseudovirus infectivity and syncytia formation in lung cells is inhibited by nitazoxanide. (**A-D**) HEK293T cells transfected with plasmids for production of SARS-CoV-2 Alpha, Beta, Gamma and Delta spike pseudotyped-LUC LV particles, as described in Fig. 4A, were treated with 25 µg/ml of NTZ or vehicle for 48h. Caco-2 hACE2 cells were infected with SARS-CoV-2 spike-pseudotyped LV-particles obtained as described above and pseudovirus entry was analyzed by measuring LUC activities 72h post infection. Data are expressed as percent of relative untreated controls. Error bars indicate means±SD. **P<0.01; Student’s *t*-test. (**E-H**) HEK293T cells co-transfected with plasmids encoding SARS-CoV-2 Alpha, Beta, Gamma or Delta S variants together with GFP for 4h and treated with NTZ (5 µg/ml) or vehicle (Control) for 36h were overlaid on A549 hACE2 cell monolayers. After 4h, cell-cell fusion was assessed by fluorescence microscopy (left panel). Bright field (BF) and merge images are shown. Scale bar, 200 µm. Cell-cell fusion was determined and expressed as percentage relative to control (right panel). Error bars indicate means±SD. *P<0.05; Student’s *t*-test. (**I**) Schematic representation of the effect of nitazoxanide on SARS-CoV-2 spike variants maturation, syncytia formation (Cell-cell fusion) and infectivity (Cell entry). Red dots depict immature forms of the spike protein (IM-Spike).

Next, we investigated the ability of nitazoxanide to prevent syncytia formation induced by the Alpha, Beta, Gamma and Delta variants. HEK293T cells were co-transfected with the plasmids of interest and GFP and, after 4h, treated with nitazoxanide or vehicle. At 40h post transfection cells were overlaid on target A549 hACE2 cells monolayers, and processed for fluorescence microscopy to visualize syncytia formation, as described above. As shown in Fig. 8E-I, all spike variants were able to cause cell-cell fusion, and treatment with nitazoxanide markedly decreased the number and size of syncytia in lung cells. As shown above, NTZ-induced inhibition of syncytia formation was dose-dependent (Fig. S8).

Altogether these results indicate that nitazoxanide is effective in inhibiting cell-cell fusion and syncytia formation mediated by different SARS-CoV-2 spike variants and in different types of cells.

## DISCUSSION

As of 22 December 2021, SARS-CoV-2 virus infections have caused more than 277 million confirmed cases and over 5.3 million deaths (https://coronavirus.jhu.edu/map.html) worldwide, wreaking havoc on global public health and economy. Given the proportion of the COVID-19 pandemic and the rise in the associated global death toll, major efforts have been directed over the past months towards a global vaccination plan (https://cdn.who.int/media/docs/default-source/immunization/covid-19/strategy-to-achieve-global-covid-19-vaccination-by-mid-2022.pdf?sfvrsn=5a68433c_5). At the same time, the emergence of several SARS-CoV-2 spike variants that facilitate virus spread and may affect the efficacy of recently developed vaccines [20–23], together with the short-lasting protective immunity typical of HCoV [65], creates great concern and highlights the importance of identifying antiviral drugs to reduce SARS-CoV-2-related morbidity and mortality. Considerable efforts have been directed over the past months towards the concept of repurposing FDA-approved drugs that have the potential to greatly accelerate clinical availability by eliminating extensive safety testing required for novel drugs. So far, only the RNA-dependent RNA polymerase (RdRp) inhibitor remdesivir has been approved by health authorities for coronavirus infections [66], whereas a different RdRp inhibitor, molnupinavir [67], was approved by MHRA (Healthcare products Regulatory Agency) in the UK (https://www.gov.uk/government/news/first-oral-antiviral-for-covid-19-lagevrio-molnupiravir-approved-by-mhra). Very recently the U.S. Food and Drug Administration also issued an emergency use authorization for Paxlovid (SARS-CoV-2 3CL protease inhibitor co-packaged with ritonavir for oral use) (https://www.fda.gov/news-events/press-announcements/coronavirus-covid-19-update-fda-authorizes-first-oral-antiviral-treatment-covid-19).

Nitazoxanide has been used for decades in medical practice as a safe and effective antiprotozoal drug [24, 25]. As described above, nitazoxanide has been proven to have a broad-spectrum antiviral activity in laboratory settings, as well as in clinical studies that have shown that the drug is effective against rotavirus, hepatitis C, and influenza A and B virus infections [26–34].

In the case of *Coronaviridae*, nitazoxanide is effective against several animal and human strains, including the MERS-CoV in cell culture [26,35,36]. As for SARS-CoV-2, at an early stage of the pandemic, Wang et al. reported that nitazoxanide inhibits SARS-CoV-2 replication in Vero E6 cells at low-μM concentrations (EC_50_  =  2.12  µM) [37]; these observations were recently confirmed in different types of cells, including human lung-derived Calu-3 cells [38, 39]. More importantly several studies have recently shown an antiviral activity and clinical benefits of nitazoxanide in COVID-19 patients [39–42].

While the molecular mechanism of its broad-spectrum antiviral activity has not been fully elucidated, nitazoxanide acts through a cell-mediated mechanism [26, 27] and, in the case of influenza and parainfluenza viruses, it was shown to impair terminal glycosylation and intracellular trafficking of the class-I viral fusion glycoproteins influenza hemagglutinin and paramyxovirus fusion spike proteins [28, 29]. This effect has been previously associated to the drug-mediated inhibition of ERp57, an ER-resident glycoprotein-specific thiol-oxidoreductase which is essential for correct disulfide-bond architecture of selected viral proteins [29].

We now show that nitazoxanide selectively interferes with SARS-CoV-2 S-glycoprotein biosynthesis, while it does not affect the expression of the other SARS-CoV-2 structural proteins N, E and M. We find that, as in the case of influenza and parainfluenza fusion proteins, nitazoxanide hampers the spike protein maturation at an Endo-H-sensitive stage, thus preventing its final processing, and resulting in inhibition of the S-glycoprotein fusion activity during entry, as well as of syncytia-formation due to S-driven cell-cell fusion. The bioactive NTZ metabolite tizoxanide was found to be equally effective in preventing SARS-CoV-2 spike maturation and syncytium-forming ability, hindering viral entry in SARS-CoV-2 S-engineered pseudovirions infection models, and inhibiting infectious SARS-CoV-2 virus propagation in Vero E6 cells with an EC_50_ of 2.9 μM.

Since its emergence, the ancestral Wuhan strain has evolved rapidly, originating several variants harboring multiple mutations, including mutations in the spike protein, which may result in the ability to evade part of the neutralizing antibody response [21–23]. We found that nitazoxanide ability to interfere with the S-glycoprotein maturation is independent of the spike variants, being equally effective against the original Wuhan-spike as well as the Alpha, Beta, Gamma and Delta spike-variants in human cells. This is in line with the cell-mediated mechanism of action previously reported for NTZ [26–29], and represents an advantage as compared to virus-targeted drugs, considering the continuous emergence of SARS-CoV-2 variants and the CoV ability to undergo recombination events [63]. Interestingly, NTZ was also found to interfere with the maturation of the spike proteins of the human alpha coronavirus 229E and beta coronavirus OC43, causing a remarkable decrease in infectious progeny production in human cells (Piacentini S. et al., manuscript in preparation), and confirming a general effect of the drug against diverse human coronaviruses.

The ability of SARS-CoV-2 S protein to mediate fusion between the viral particle and the cellular membranes during virus entry has been largely studied and documented; however, there is recent evidence that the spike protein is also responsible for fusion of infected cells with neighboring uninfected cells [59, 68]. This process, known as ‘fusion from within’, is mediated by newly synthesized S protein reaching the cell surface, fusing with receptor-positive adjacent cells, and causing the formation of giant multinucleated cells (syncytia). Cell-cell fusion is used by different viruses, including the Pneumovirus RSV (Respiratory Syncytial virus), the measles virus (MV) [69] and the human immunodeficiency virus (HIV) [70], to spread in a particle-independent way and promoting immune evasion [71].

Utilizing two different quantitative assays relying on the biogenesis of expressed S protein, we now confirm that the SARS-CoV-2 Wuhan-spike as well as the Alpha, Beta, Gamma and Delta spike-variants possess remarkable cell-cell membrane fusion activity, causing syncytia formation in ACE2-expressing cells, and demonstrate that nitazoxanide is effective in inhibiting syncytia formation mediated by the different spike-variants in different types of cells.

Syncytia formation has been linked to pathological consequence detectable in various tissues such as lung and brain for MV [72] or lymphoid tissues for HIV [70]. In the case of SARS-CoV-2 infection, syncytia formation has been recently described as an hallmark of advanced lung pathology in patients affected with COVID-19 in an occurrence not seen in other lung infections before [58,73–76]. In particular, fused pneumocytes expressing SARS-CoV-2 RNA and S proteins were observed post-mortem in lung tissues of 20 out of 41 COVID-19-infected patients, indicating that productive infection leads to syncytia formation [58]. Notably SARS-CoV-2 S-mediated syncytia were suggested to increase viral pathogenicity by destabilizing airway epithelia and creating lesions that may be difficult to repair [75]. Interestingly, it has also been recently shown that, whereas neutralizing antibodies and sera from convalescent patients inhibited SARS-CoV-2 particle-cell fusion with high efficiency, cell-cell fusion was only moderately inhibited, suggesting that syncytia formation during COVID-19 may not be effectively prevented by antibodies [68].

The results described herein suggest that, together with virus-targeting drugs, the host-directed broad-spectrum antiviral drug nitazoxanide, by hampering the spike protein maturation and fusion activity, may represent a readily-available useful tool in the fight against COVID-19 infections, inhibiting SARS-CoV-2 replication and preventing spike-driven syncytia formation.

## Materials and Methods

### Cell culture, treatments, plasmids and transfections

Human alveolar type II-like epithelial A549 cells, airway epithelial Calu-3 cells and MRC-5 lung fibroblasts, colorectal adenocarcinoma HCT116 cells, colon carcinoma Caco-2 cells, embryonic kidney 293T cells, and African green monkey kidney Vero E6 cells, murine L929 fibroblasts and bat lung epithelial Tb-1 Lu cells were obtained from ATCC (Manassas, VA, USA). The human Huh7 hepatoma cell line was kindly provided by Massimo Levrero (INSERM, Lyon, France). Cells were grown at 37°C in a 5% CO_2_ atmosphere in RPMI-1640 (Euroclone; A549 cells), DMEM (Euroclone; Vero E6, Huh7, 293T, HTC116 and Caco-2 cells), EMEM (ATCC; MRC-5 and Calu-3 cells) (Lonza; Tb-1 Lu cells) or MEM (Gibco; L929 cells) medium supplemented with 10% fetal calf serum (FCS), 2 mM glutamine and antibiotics. Cell viability was determined by the 3-(4,5-dimethylthiazol-2-yl)-2,5-diphenyltetrazolium bromide (MTT) to MTT formazan conversion assay (Sigma-Aldrich) as described [29].

Nitazoxanide [2-acetyloxy-N-(5-nitro-2-thiazolyl) benzamide, Alinia] and tizoxanide (Romark Laboratories), remdesivir (RDV, MedChemExpress), bortezomib (BTZ, Selleckchem), kifunensine (Cayman), ribavirin (RBV), chloroquine (CQ), cycloheximide (CHX), decanoyl-RVKR-CMK (RVKR), tunicamycin (TM), brefeldin A (BFA), castanospermine (CST) and 1-deoxymannojirimicin (DMJ) (Sigma-Aldrich), dissolved in DMSO stock solution (NTZ, TIZ, RDV, RVKR, BTZ, kifunensine, BFA) or aqueous solution (RBV, CQ, CHX, TM, CST, DMJ), were diluted in culture medium and maintained in the medium for the duration of the experiment. Controls received equal amounts of vehicle.

The pCMV3-2019-nCoV-Spike(S1+S2)-long (SARS-2 S) construct and the C-terminal Flag-tagged pCMV3-2019-nCoV-Spike(S1+S2)-long-Flag (SARS-2 S-CF), pCMV3-SARS-spike-Flag (SARS S) and pCMV3-Spike (betacoronavirus-2c-EMC2012)-Flag (MERS S), pCMV3-SARS-CoV-2-Spike (S1+S2)-long(B.1.617.2) (Delta), pCMV3-2019-nCoV-M-Flag (SARS-2 M), pCMV3-2019-nCoV-ENV-Flag (SARS-2 E) and the C-terminal c-Myc-tagged pCMV3-2019-nCoV-NP-Myc (SARS-2 N) vectors were obtained from Sino Biological. The SARS-CoV-2 variants C-terminal Flag-tagged pReceiver-M39 U.K. B.1.1.7 (EX-CoV242-M39, Alpha), South Africa B.1.351 (EX-CoV244-M39-GS, Beta) and Brazil B.1.1.28.1/P.1 (EX-CoV245-M39-GS, Gamma) vectors were obtained from GeneCopoeia. The mutants N501Y, N501Y+E484K and K417N+E484K+N501Y were constructed on the SARS-2 S-CF plasmid; mutations were generated and sequence-validated by Bio-Fab Research (Rome, IT). C-terminal Flag-tagged pCAGGS-SARS2-S-D614 (Addgene plasmid #156420) and pCAGGS-SARS2-S-G614 (Addgene plasmid #156421) vectors were a gift from Hyeryun Choe and Michael Farzan [77]. The pCMV-GFP vector was obtained from Clontech. Transfections were performed using jetPRIME Transfection-Reagent (Polyplus-transfection), according to the manufacturer’s instructions.

### Generation of A549 hACE2, 293T hACE2 and Caco-2 hACE2 stable cell lines

Lentiviral particles to generate hACE2 stably expressing cell lines were produced by co-transfection of 293T cells with the VSV-G encoding plasmid [78], the pCMVR 8.74 lentiviral packaging plasmid (a kind gift from R. Piva, University of Turin, Italy) and the pLENTI hACE2 PURO vector (a gift from Raffaele De Francesco, INGM, Milan; Addgene plasmid #155295). At 48h after transfection, supernatants were collected, filtered through 0.45-mm membranes and used to infect A549, 293T and Caco-2 cells in the presence of 8 μg/ml of polybrene, and then selected with puromycin (1 µg/ml) 72h after infection. After 10 days in selective medium resistant pools of A549, 293T and Caco-2 cells were obtained. The puromycin selective pressure was removed 24h before experimental procedures.

### Protein Analysis, Western blot and endoglycosidase digestion

For analysis of soluble/insoluble proteins whole-cell extracts (WCE) were prepared after lysis in High Salt Buffer (HSB) [79] without DTT. Briefly, cells were washed twice with ice-cold PBS and then lysed in HSB (80µl). After one cycle of freeze and thaw, and centrifugation at 16,000×g (10min at 4°C), supernatant (soluble) and pellet (insoluble) fractions were collected. Insoluble fractions were solubilized in 60µl of Buffer-S (50mM Tris-HCl, pH 8.5, 1% SDS and protease inhibitors) by sonication on ice, using an ultrasonic UP50H processor (Hielscher) (40% amplitude, pulse mode: 6×10 sec, 15 sec pauses) [79].

For Western blot analysis, cell extracts (20 µg/sample) were separated by SDS-PAGE under non-reducing conditions and blotted to a nitrocellulose membrane. Membranes were incubated with the selected antibodies, followed by incubation with peroxidase-labeled anti-rabbit or anti-mouse IgG. Primary and secondary antibodies used are listed in Supplementary Table 1.

For endoglycosidase digestion experiments, samples containing the same amount of protein (50 µg/sample) were processed for endoglycosidase-H (Endo-H, NEB) digestion using 5 milliunits Endo-H for 16h at 37°C, or for peptide N-glycosidase F (PNGase-F, NEB) digestion using 500 units PNGase-F for 1,5h at 37°C, according to the manufacturer’s protocol [28]. Samples were separated by SDS-PAGE under reducing conditions and blotted as described above. Quantitative evaluation of proteins was determined as described [80–82]. All results shown are representative of at least three independent experiments.

### Immunofluorescence microscopy

A549 cells transfected with Flag-tagged SARS-CoV-2 spike or empty vector were grown in 8-well chamber slides (Lab-Tek II) and, after 4h, were treated with NTZ, TM or vehicle for 16h. Cells were fixed, permeabilized and processed for immunofluorescence as described [83] using anti-Flag antibodies, followed by decoration with Alexa Fluor 488- or 555-conjugated antibodies (Molecular Probes, Invitrogen). Nuclei were stained with Hoechst 33342 (Molecular Probes, Invitrogen). Images were captured using a ZEISS Axio Observer Inverted Microscope and analyzed using ZEN 3.1 (blue edition) software. For confocal microscopy, images were acquired on Olympus FluoView FV-1000 confocal laser scanning system (Olympus America Inc., Center Valley, PA) and analyzed using Imaris (v6.2) software (Bitplane, Zurich, Switzerland). Images shown in all figures are representative of at least three random fields (scale-bars are indicated).

### Cell-cell fusion assay

Two different protocols were used for visualization of syncytia. In the first protocol, Vero E6, Huh7, A549 and A549 hACE2 cells grown in 8-well chamber slides were transfected with Flag-tagged SARS-CoV-2 (SARS-2 S-CF wild type or mutated, pCAGGS-SARS2-S-D614, pCAGGS-SARS2-S-G614) or SARS-CoV spike plasmids or empty vector and, after 4h, were treated with NTZ or vehicle. After 36h, cells were processed for immunfluorescence as described above using an anti-Flag antibody. Nuclei were stained with Hoechst. Syncytia images were captured using a ZEISS Axio Observer inverted microscope and analyzed using ZEN 3.1 (blue edition) software. For syncytia formation quantification, the percentage of cells involved in syncytia formation was determined by counting the number of cells containing 3 or more nuclei, compared with the number of total nuclei in transfected cells in the same sample [57]. At least 1.000 nuclei were counted for each sample.

In the donor-target cell fusion assay, 293T cells were co-transfected with plasmids encoding Flag-tagged SARS-CoV-2 S-glycoprotein alone or in combination with Flag-tagged SARS-CoV-2 M-glycoprotein and E protein, and c-Myc-tagged N protein together with GFP using Lipofectamine 2000 (Life Technologies), according to the manufacturer’s instructions. For SARS-CoV-2 variants cell-fusion assay, 293T cells were co-transfected with the plasmids encoding SARS-CoV-2 S-glycoprotein pReceiver-M39 U.K. B.1.1.7 (EX-CoV242-M39) (Alpha), South Africa B.1.351 (EX-CoV244-M39-GS) (Beta), Brazil B.1.1.28.1/P.1 (EX-CoV245-M39-GS) (Gamma) or pCMV3-SARS-CoV-2-Spike (S1+S2)-long(B.1.617.2) (Delta) together with GFP, as described above. After 4h, cells were treated with NTZ or vehicle. 293T cells transfected with the GFP-encoding plasmid were used as negative control. Cells were detached with trypsin (0.25%) at 40h post transfection and overlaid on an 80% confluent monolayer of target cells [Vero E6 and Huh7 (naturally expressing hACE2 receptors on the membrane surface) or A549-hACE2 (stably transfected with the hACE2 encoding plasmid) cells] at a ratio of approximately one S-expressing cell to three receptor-expressing cells [45]. After 4h, nuclei were stained with Hoechst. Transmission and fluorescence images were taken using a ZEISS Axio Observer inverted microscope. To measure the extent of cell–cell fusion, the GFP area was delimited, measured using ImageJ software [59], divided by the total cell area and expressed as percentage relative to control.

Images shown in all figures are representative of at least five random fields (scale-bars are indicated). All experiments were done in duplicate and repeated at least three times.

### Generation of SARS-CoV-2 S-pseudoviruses

For production of Murine Leukemia Virus (MLV)-based SARS-CoV-2 S-pseudotyped particles, 293T cells grown in 60mm dishes were co-transfected with the SARS-2 S-CF plasmid encoding the SARS-CoV-2 S-protein, the MLV Gag-Pol packaging construct [78] and the MLV transfer vector encoding a luciferase reporter (PTG-Luciferase) [78]. For production of lentiviral SARS-CoV-2 S-pseudotyped particles, 293T cells were co-transfected with the SARS-2 S-CF plasmid [or alternatively, the pCAGGS-SARS2-S-D614, pCAGGS-SARS2-S-G614, pReceiver-M39 U.K. B.1.1.7 (EX-CoV242-M39) (Alpha), South Africa B.1.351 (EX-CoV244-M39-GS) (Beta), Brazil B.1.1.28.1/P.1 (EX-CoV245-M39-GS) (Gamma) or pCMV3-SARS-CoV-2-Spike (S1+S2)- long(B.1.617.2) (Delta) plasmids], the pCMVR 8.74 lentiviral packaging plasmid and the pLenti CMV Puro LUC or pLenti CMV Puro GFP vectors. pLenti CMV Puro LUC (Addgene plasmid #17477) and pLenti CMV Puro GFP (Addgene plasmid #17448) vectors were a gift from Eric Campeau and Paul Kaufman [84]. Transfections were performed using Lipofectamine 2000 (Life Technologies), according to the manufacturer’s instructions. After 5h, cells were washed twice with medium and treated with different concentrations of NTZ or vehicle. At 48h after transfection, supernatants containing the viral particles were collected and filtered through 0.45 mm membranes. Pseudoviruses were purified using PEG Virus Precipitation Kit (Abcam), resuspended in 100 μl of virus re-suspension solution and then frozen at -80°C until use. After collection of viral particles, the monolayers of 293T producing cells were lysed and analyzed for S-protein levels as described above.

### Pseudovirus entry assays

Vero E6, Calu-3, Caco-2, Caco-2 hACE2 and 293T hACE2 cells were plated at a density of 2x10^4^ cells/well in 96 well white clear-bottom plates (Costar). After 18h, 0.1 ml of DMEM containing 5% FCS and 8 µg/ml polybrene was added to each well and cells were incubated for 30min. The medium was then removed and the pseudovirus suspension (50 μl: 5 μl of purified pseudovirus stock + 45 μl of DMEM containing 5% FCS) was added; after 24h, 0.05 ml of DMEM 5% FCS was added to each well. Luciferase activity (Relative Luciferase Units or RLU) was detected at 72h post-infection by using Bright-Glo Luciferase Assay System Kit (Promega) in a Microplate Luminometer (Wallac-Perkin Elmer).

All experiments were performed in triplicates, and repeated at least three times.

### Antiviral activity assay

To determine the antiviral activity of tizoxanide, Vero E6 cells were infected with SARS-CoV-2 (SARS-CoV-2/England/2/2020) virus for 2h, and then treated with a three-fold serial dilution of compound ranging from 20 µM to 0.01 µM. Antiviral activity was determined at 24h using an immunofluorescence-based assay. Briefly, cells were fixed, permeabilized and stained with an antibody recognizing SARS-CoV-2 spike protein (GeneTex GTX632604). The primary antibody was detected with an Alexa-488 conjugate secondary antibody (Life Technologies, A11001), and nuclei were stained with Hoechst. Images were acquired on an Opera Phenix high content confocal microscope (Perkin Elmer) using a 10X objective (9 images per well, covering approx. 2/3 of the well), and percentages of infection were calculated using Columbus software (infected cells/total cells x 100). The percentage of inhibition of SARS-CoV-2 infection was normalized to the internal controls (100% inhibition is derived from the average of the untreated uninfected control and 0% inhibition is derived from the average of the untreated infected control). Cell viability was determined by MTT assay [29]. Cytotoxicity values were normalized to the internal control (untreated cells, 100% viability). Tizoxanide antiviral activity assays against SARS-CoV-2 were performed by Virology Research Services Ltd (UK).

### Statistical analysis

Statistical analysis was performed using the Student’s *t-*test for unpaired data or one-way ANOVA test (Prism 5.0 software, GraphPad). Data are expressed as the mean±SD of samples derived from at least three biological repeats and *P* values ≤0.05 were considered significant. All the results shown are representative of at least three independent experiments, each in duplicate or triplicate.

### Author Contributions

A Riccio, SS and SP performed the analysis of protein synthesis, maturation and intracellular localization; A. Rossi performed the pseudovirus studies; A. Riccio and SS conducted the cell-cell fusion study; MGS and JFR designed the study; MGS, AR and SS wrote the manuscript. All authors contributed to the interpretation of the data and approve the content of the manuscript.

## Supporting information

Riccio et al, Supplementary Information

## Acknowledgments

This research was supported by Romark Laboratories LC, Tampa, Florida, and by a grant from the Italian Ministry of University and Scientific Research (PRIN project N 2010PHT9NF-006). The authors thank Massimo Levrero (INSERM U1052, Lyon, France) for providing the Huh7 hepatoma cell line.

