## Supplementary material for "Impairment of SARS-CoV-2 spike-glycoprotein maturation and fusion-activity by nitazoxanide: an effect independent of spike variants emergence": Riccio et al, Supplementary Information

**Supplementary Table 1. Antibodies used.**

| Antibody | Source | Catalogue Number |
|---|---|---|
| Calnexin (M) | Thermo Fisher Scientific (IF) | MA3-027 |
| KDEL (M) | Enzo Life Sciences | ADI-SPA-827 |
| SARS-CoV-2 spike (M) | Sino Biological | 40150-R007 |
| α-Tubulin (M) | Sigma-Aldrich | T5168 |
| ACE2 (M) | Santa Cruz | SC-73668 |
| β-Actin (P) | Sigma-Aldrich | A2066 |
| c-Myc (M) | Santa Cruz | SC-40 (9E10) |
| Calnexin (P) | StressGen (WB) | ADI-SPA-865 |
| Calreticulin (P) | StressGen | ADI-SPA-600 |
| DYKDDDDK Tag, Flag (P) | Cell Signaling | 2368 |
| ERp57 (P) | Merck Millipore | ABE1032 |
| ERp72 (P) | Enzo Life Sciences | ADI-SPA-720 |
| Alexa Fluor 488 goat anti-mouse | Invitrogen | A11001 |
| Alexa Fluor 555 goat anti-rabbit | Invitrogen | A21428 |
| Goat Anti-Mouse IgG (H+L), HRP | Jackson ImmunoResearch | 115-035-003 |
| Goat Anti-Rabbit IgG (H+L), HRP | Jackson ImmunoResearch | 111-035-003 |

(**M**): Monoclonal; (**P**): Polyclonal

### Supplementary Figure Legends

**Figure S1. Nitazoxanide affects SARS-CoV-2 spike biogenesis in different types of mammalian cells.**

**(A-E)** Human liver (Huh7), kidney (HEK293T) and lung (A549) cells (**A,B**), or monkey Vero E6 (**C**), murine L929 (**D**) and bat lung epithelial Tb 1 Lu (**E**) cells were transiently transfected with the C-Flag tagged SARS-CoV-2 spike (SARS-2 S-CF) construct or empty vector (Mock) and, after 4h (Huh7, HEK293T, A549, Vero E6 and L929) or 6h (Tb 1 Lu), were treated with different concentrations (**A, C-E**) or 2.5 μg/ml (**B**) of NTZ or vehicle. At different times (**B**) or at 16h (**A, C-E**) after treatment, whole cell extracts were analyzed for S protein levels by IB using anti-Flag or anti-spike antibodies. Black arrows indicate bands corresponding to uncleaved S proteins (S0), whereas gray arrows indicate bands corresponding to the S1/S2 subunits. The slower- and faster-migrating S0 and S1/S2 forms in untreated or NTZ-treated cells are identified by asterisk and arrowheads respectively. Data from a representative experiment of three with similar results are shown.

**Figure S2. Nitazoxanide inhibits the fusogenic activity of SARS-CoV-2 spike proteins carrying single or multiple mutations in human lung cells.**

(**A**) A549 cells were transiently transfected with the SARS-2 S-CF plasmid carrying the N501Y mutation (N501Y), double N501Y and E484K mutations (N501Y+E484K) or triple K417N, E484K and N501Y mutations (K417N+E484K+N501Y) or empty vector, and treated with NTZ (2.5 μg/ml) or vehicle (-). After 16h, WCE were analyzed for levels of S protein by IB using anti-Flag antibodies. Black arrows indicate bands corresponding to uncleaved S protein (S0), whereas gray arrows indicate bands corresponding to the S2 subunit. The slower- and faster-migrating S0 and S2 forms in untreated or NTZ-treated cells are identified by asterisk and arrowheads respectively. (**B**) A549 cells stably expressing human ACE2 (A549 hACE2) were transiently transfected with SARS-2 S-CF N501Y, N501Y/E484K or K417N/E484K/N501Y plasmids described in (**A**) for 4h and treated with NTZ (5 μg/ml) or vehicle (Control) for 36h. Immunofluorescence analysis was performed using an anti-Flag antibody (red) (left panels). Nuclei are stained with Hoechst (blue). Merge and zoom images are shown. Scale bar, 200 μm (zoom, 100 μm). The number of nuclei involved in syncytia formation is expressed as percent of total nuclei in transfected cells in the same sample (right panel). Data represent the means±SD of 5 fields from three biological replicates. *P<0.05; **P<0.01; Student's *t*-test. (**C**) HEK293T cells co-transfected with plasmids encoding GFP together with SARS-2 S-CF plasmids carrying the N501Y mutation (N501Y), double E484K and N501Y mutations (E484K/N501Y) or triple K417N, E484K and N501Y mutations (K417N/E484K/N501Y) for 4h and treated with NTZ (5 μg/ml) or vehicle for 36h were overlaid on A549 hACE2 cell monolayers. After 4h, cell-cell fusion was assessed by fluorescence microscopy (left panels). Bright field and merge images are shown. Scale bar, 200 μm. Cell-cell fusion was determined and expressed as percentage relative to control (right panel). Error bars indicate means±SD. *P<0.05; **P<0.01; Student's *t*-test.

**Figure S3. Tizoxanide inhibits SARS-CoV-2 replication and SARS-CoV-2 S pseudovirus infectivity.**

(**A**) Structure of tizoxanide (TIZ). (**B**) Vero E6 cells were infected with SARS-CoV-2 (SARS-CoV-2/England/2/2020) virus for 2h, and then treated with different concentrations of TIZ. Antiviral activity (black line) and cytotoxicity (blue line) were determined at 24h by immunofluorescence and MTT assay respectively, as described in 'Materials and Methods'. (**C,D**) HEK293T cells transfected with plasmids for production of SARS-CoV-2 spike-pseudotyped LUC MLV particles were treated with different concentrations of tizoxanide or control diluent. At 48h post-transfection, pseudovirus-containing supernatants were collected for pseudovirus entry assays (see panel **D**), whereas SARS-CoV-2 S expression was detected in WCE of HEK293T pseudovirus-producing cells by IB using anti-spike antibodies **(C).** Black arrow indicates the band corresponding to uncleaved S proteins (S0), whereas the gray arrow indicates the band corresponding to the S1 subunit. The slower- and faster-migrating S0 and S1 forms in untreated or tizoxanide-treated cells are identified by asterisk and arrowheads respectively. (**D**) hACE2-expressing Caco-2 cells were infected with SARS-CoV-2 spike pseudotyped MLV particles obtained as described in **(C)**, and pseudovirus entry was analyzed by measuring luciferase activities 72h post-infection. Data are expressed as percent of untreated control. Error bars indicate means±SD. *P<0.05 and **P<0.01; ANOVA.

**Figure S4. The *N*-glycosylation inhibitor tunicamycin causes SARS-CoV-2 S protein aggregate formation in lung cells.**

(**A**) IB analysis of S protein (α-spike), ubiquitin, β-actin and histone H3 levels in soluble and insoluble fractions of WCE from A549 cells transiently transfected with the SARS-CoV-2 spike (SARS-2 S) construct or empty vector (Mock) for 4h and treated with 2.5 μg/ml NTZ, 2.5 μg/ml tunicamycin (TM), 20 nM bortezomib (BTZ) or vehicle for 16h. Black arrows indicate bands corresponding to uncleaved S proteins (S0), whereas gray arrows indicate bands corresponding to the S1 subunits. The slower- and faster-migrating forms of the uncleaved S protein or cleaved subunit are identified by asterisks and arrowheads respectively. Red arrow indicates bands corresponding to the non-glycosylated S protein detected in the insoluble fraction of TM-treated cells. (**B**) Confocal images of Flag-tagged S protein (red) intracellular distribution in A549 cells transiently transfected with the SARS-2 S-CF construct or empty vector for 4h and treated with TM (5 μg/ml) or vehicle for 16h. Nuclei are stained with Hoechst (blue). Merge and zoom images are shown. Scale bar, 20 μm (zoom, 8 μm). Data from a representative experiment of three with similar results are shown. (**C,D**) IB analysis of S protein (α-spike) and β-actin levels of WCE from A549 cells transiently transfected with SARS-2 S (**C**) or SARS-2 S CF construct (**D**) or empty vector (Mock) for 4h and treated with 2.5 μg/ml NTZ, 2 μg/ml Brefeldin A (BFA), 50 μg/ml castanospermine (CST), 15 μg/ml 1-deoxymannojirimicin (DMJ), kifunensine (7.5 or 15 μM) or vehicle for 16h.

**Figure S5. Nitazoxanide-induced impairment of spike glycoprotein maturation and fusion activity is not affected by the presence of SARS-CoV-2 M, E and N structural proteins.**

(**A**) A549 cells were transiently transfected with C-terminal Flag tagged SARS-CoV-2 spike (SARS-2 S-CF), membrane glycoprotein (SARS-2 M), envelope protein (SARS-2 E) or C-terminal c-Myc tagged SARS-CoV-2 nucleoprotein (SARS-2 N) constructs for 4h and treated with NTZ (2.5 μg/ml) or vehicle (-). After 16h, whole cell extracts were analyzed for levels of S, N, M and E proteins by IB using anti-spike, anti-Myc or anti-Flag antibodies. (**B**) A549 were transiently co-transfected with the same amount of the SARS-2 S-CF, SARS-2 M, SARS-2 E and SARS-2 N constructs for 4h and treated with NTZ (2.5 μg/ml) or vehicle (-) for 16h. S, N, M and E protein levels were determined by IB as described in (**A**). Black arrows indicate bands corresponding to uncleaved S protein (S0), whereas gray arrows indicate bands corresponding to the S1 subunit. (**C**) HEK293T cells co-transfected with plasmids encoding SARS-CoV-2 S, N, M and E proteins and GFP for 4h and treated with NTZ (5 μg/ml) or vehicle (Control) for 36h were overlaid on A549 hACE2 cell monolayers. After 4h, cell-cell fusion was assessed by fluorescence microscopy. Bright field (BF) and merge images are shown (left panel). Scale bar, 200 μm. Cell-cell fusion was determined and expressed as percentage relative to control (right panel). Error bars indicate means±SD. $^{*}P<0.05$; Student's *t*-test.

**Figure S6. Nitazoxanide inhibits SARS-CoV-2 S pseudovirus infectivity in intestinal epithelial Caco-2 hACE2 cells.**

(**A**) SARS-CoV-2 spike pseudotyped-GFP lentiviral particles produced as described in Figure 4 were used to infect Caco-2 hACE2 cells. Pseudovirus-infected cells were visualized by fluorescence microscopy. Bright field (BF) and merge images are shown. Scale bar, 200 μm. hACE2 protein levels in Caco-2 WT and Caco-2 hACE2 cells are shown in (**B**).

**Figure S7. Nitazoxanide treatment hinders SARS-CoV-2 S-mediated cell fusion in human liver Huh7 cells.**

Syncytia formation by Huh7 cells transiently transfected with the C-Flag tagged SARS-CoV-2 spike construct for 4h and treated with NTZ (5 μg/ml) or vehicle (Control) for 36h were analyzed by immunofluorescence microscopy (left panel) using an anti-Flag antibody (red). Nuclei are stained with Hoechst (blue). Merge images are shown. Scale bar, 50 μm. The number of nuclei involved in syncytia formation is expressed as percent of total nuclei in transfected cells in the same sample (right panel). Data represent the mean±SD of 5 fields from three biological replicates. *P<0.05; Student's *t*-test.

**Figure S8. Dose-dependent inhibition of SARS-CoV-2 spike-driven lung cell syncytia formation by nitazoxanide.**

(**A,B**) HEK293T cells co-transfected with plasmids encoding SARS-CoV-2 Wuhan (**A**) or Delta (**B**) spike variants together with GFP for 4h and treated with different concentrations of NTZ or vehicle (Control) for 36h were overlaid on A549 hACE2 cell monolayers. After 4h, cell-cell fusion was assessed by fluorescence microscopy (left panels). Bright field (BF) and merge images are shown. Scale bar, 200 μm. Cell-cell fusion was determined and expressed as percentage relative to control (right panels). Error bars indicate means±SD. *P<0.05, **P<0.01; ANOVA.

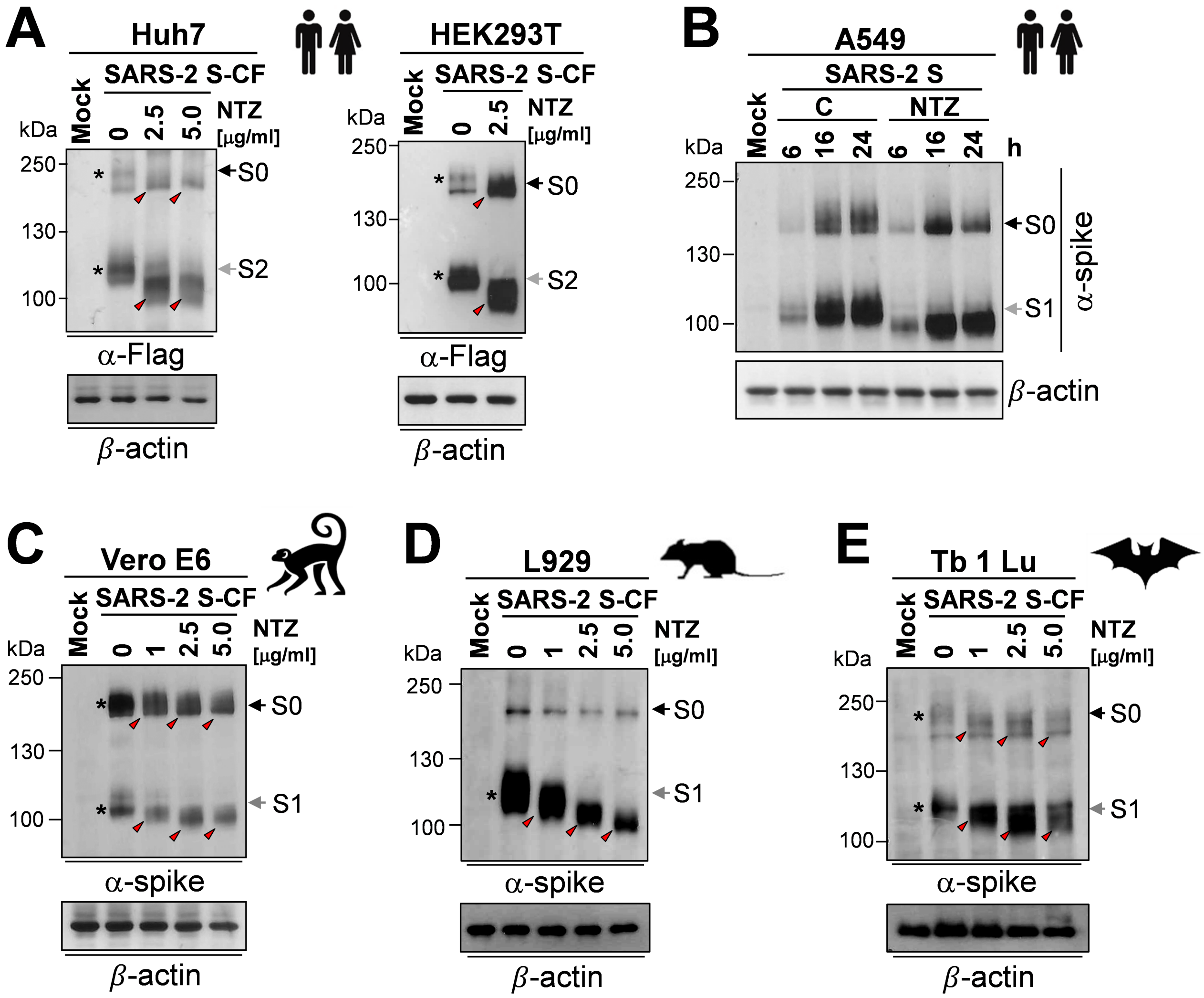


**Figure S1. Nitazoxanide affects SARS-CoV-2 spike biogenesis in different types of mammalian cells.**

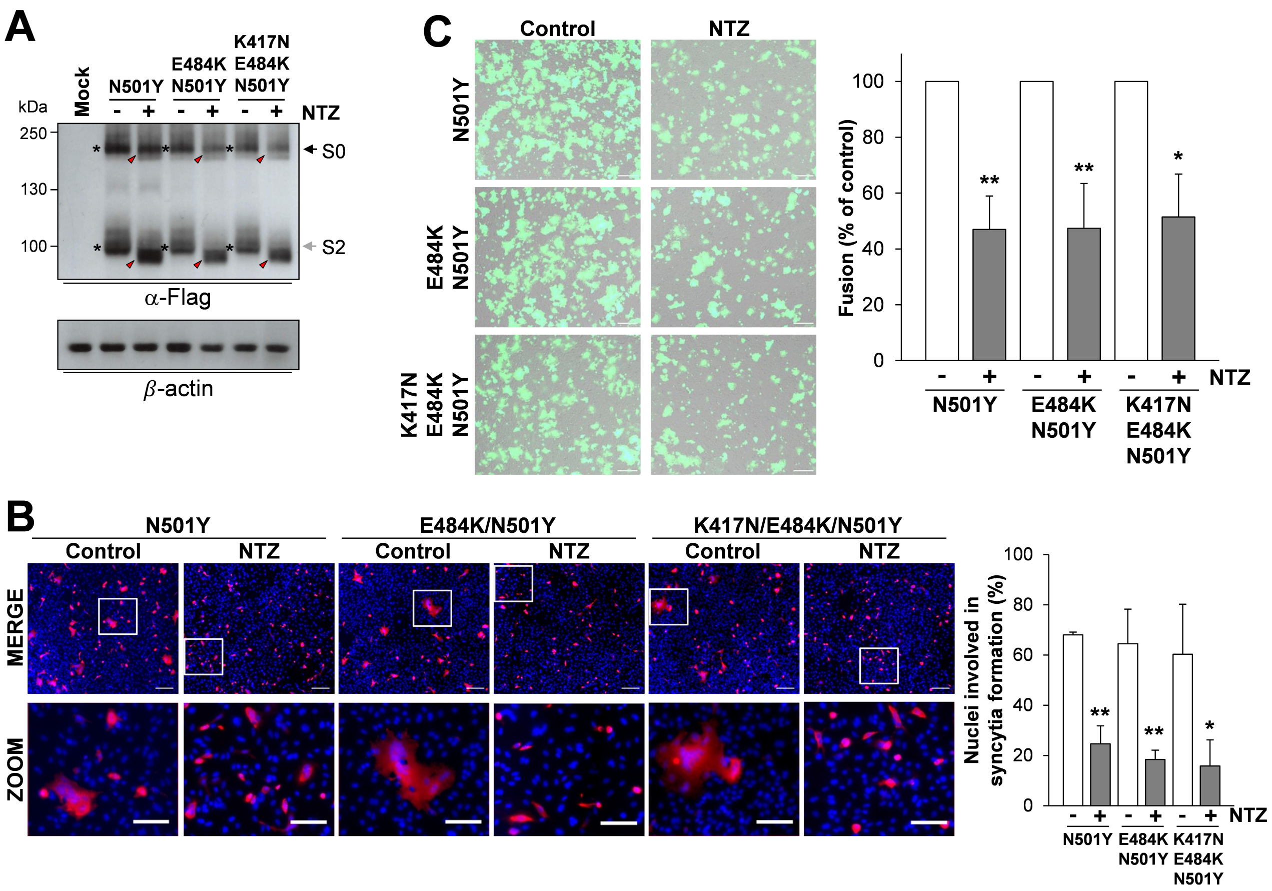


**Figure S2. Nitazoxanide inhibits the fusogenic activity of SARS-CoV-2 spike proteins carrying single or multiple mutations in human lung cells.**

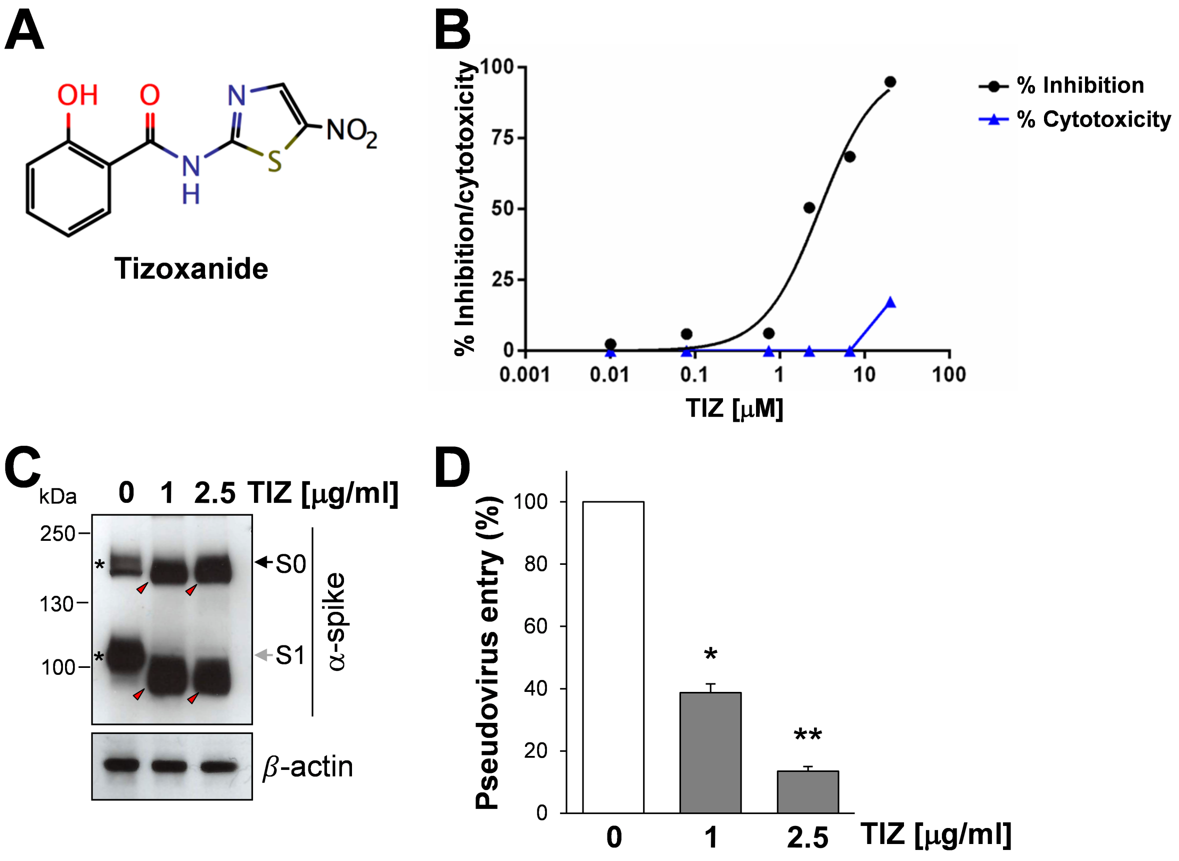


**Figure S3. Tizoxanide inhibits SARS-CoV-2 replication and SARS-CoV-2 S pseudovirus infectivity.**

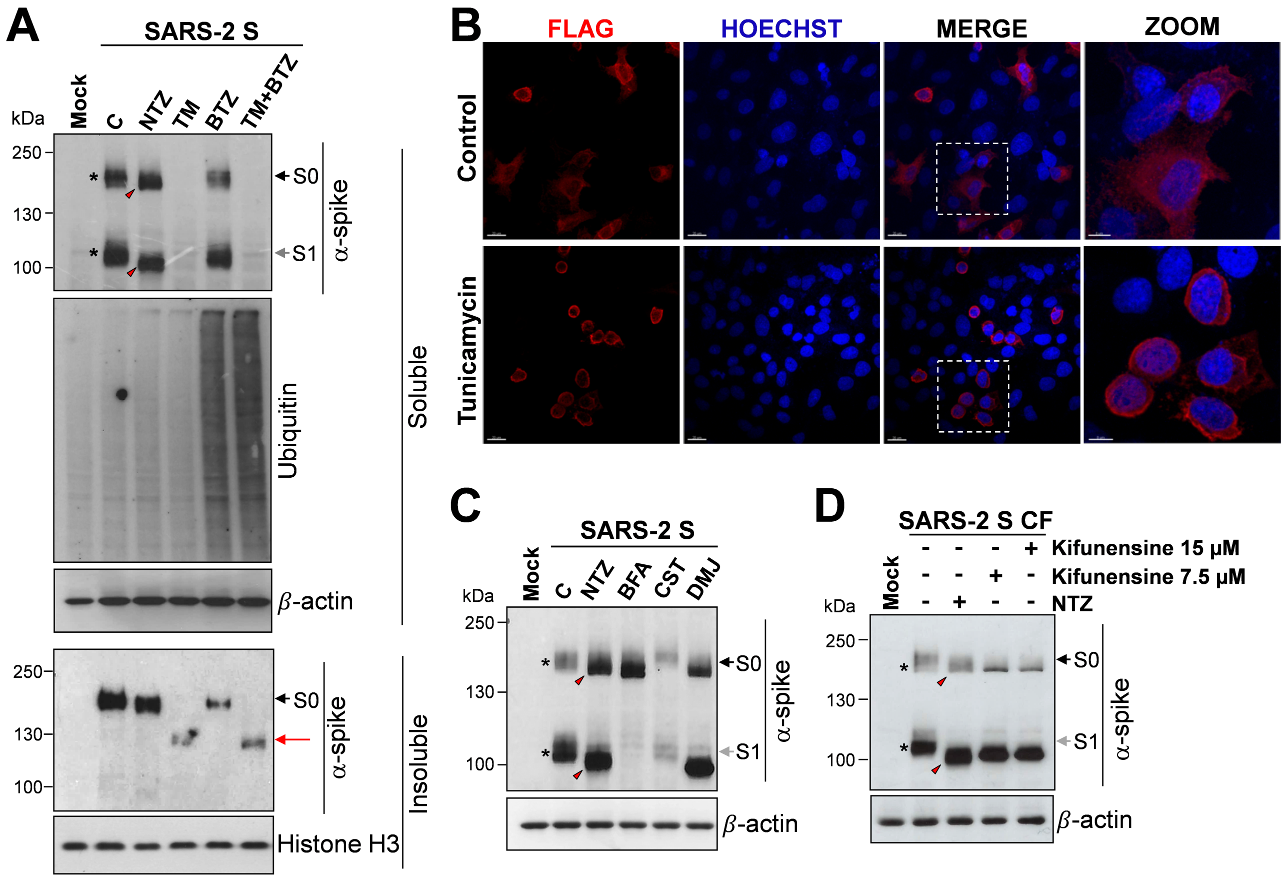


**Figure S4. The *N*-glycosylation inhibitor tunicamycin causes SARS-CoV-2 spike protein aggregate formation in lung cells.**

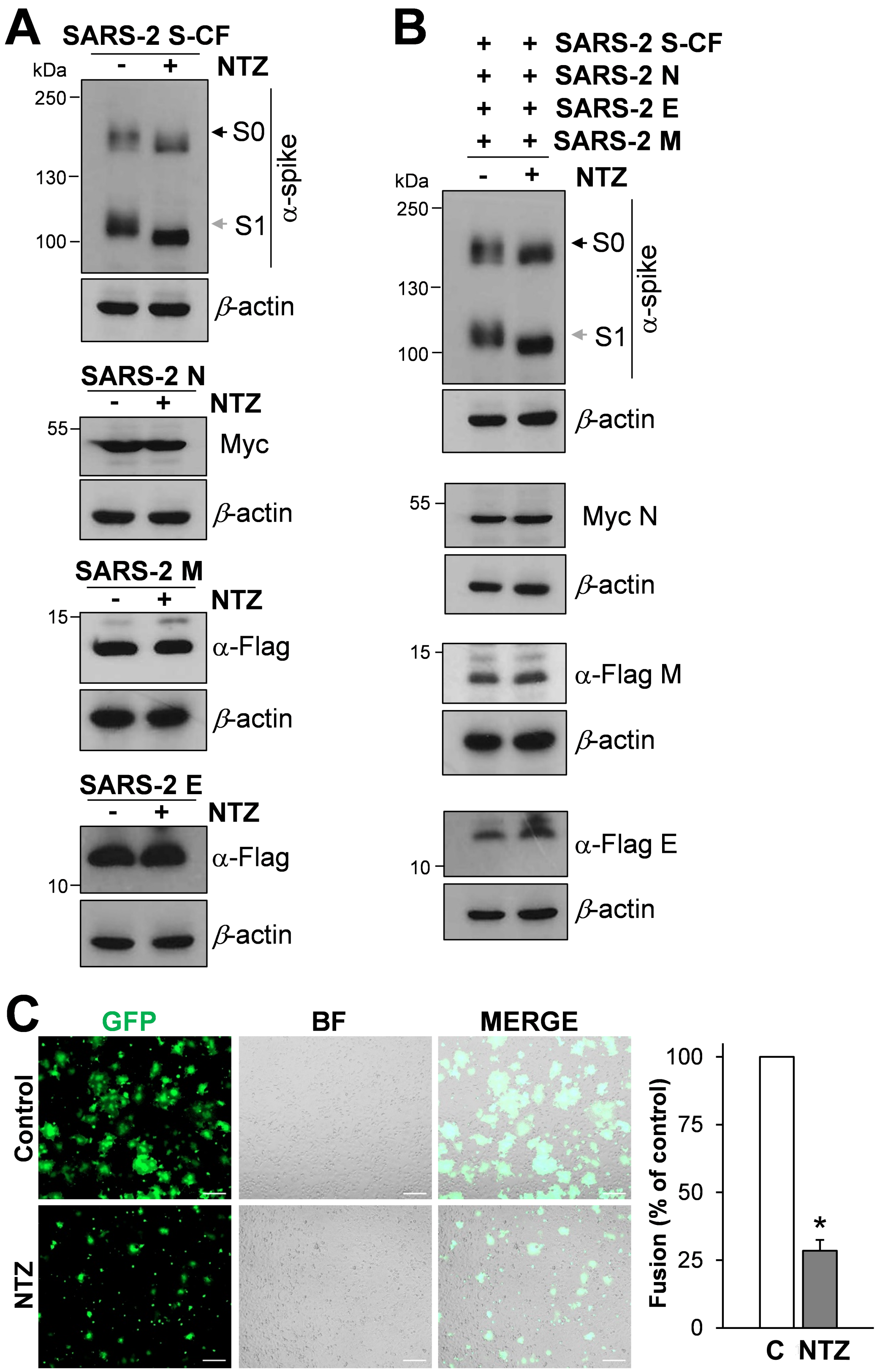


**Figure S5. Nitazoxanide-induced impairment of spike glycoprotein maturation and fusion activity is not affected by the presence of SARS-CoV-2 M, E and N structural proteins.**

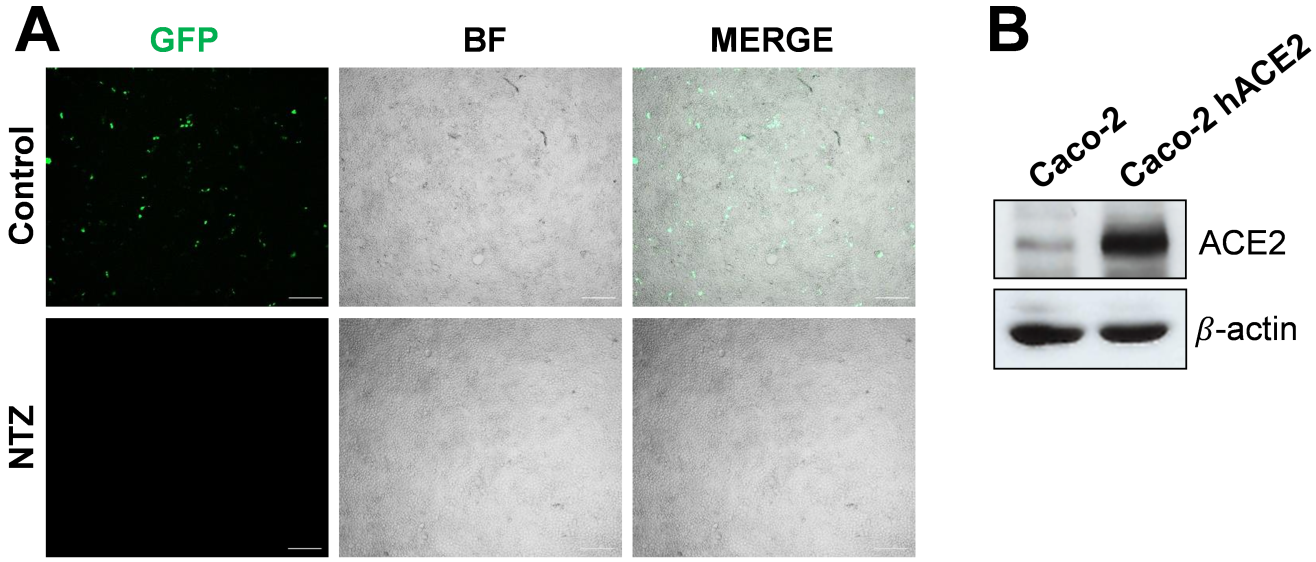


**Figure S6. Nitazoxanide inhibits SARS-CoV-2 S pseudovirus infectivity in intestinal epithelial Caco-2 hACE2 cells.**

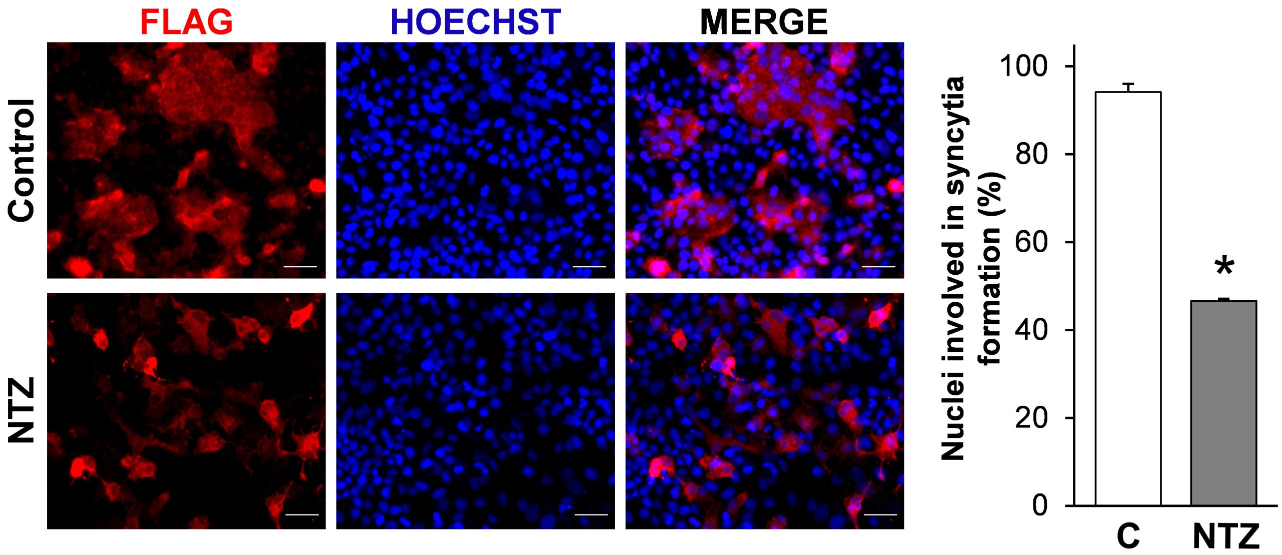


**Figure S7. Nitazoxanide treatment hinders SARS-CoV-2 S-mediated cell-cell fusion in human liver Huh7 cells.**

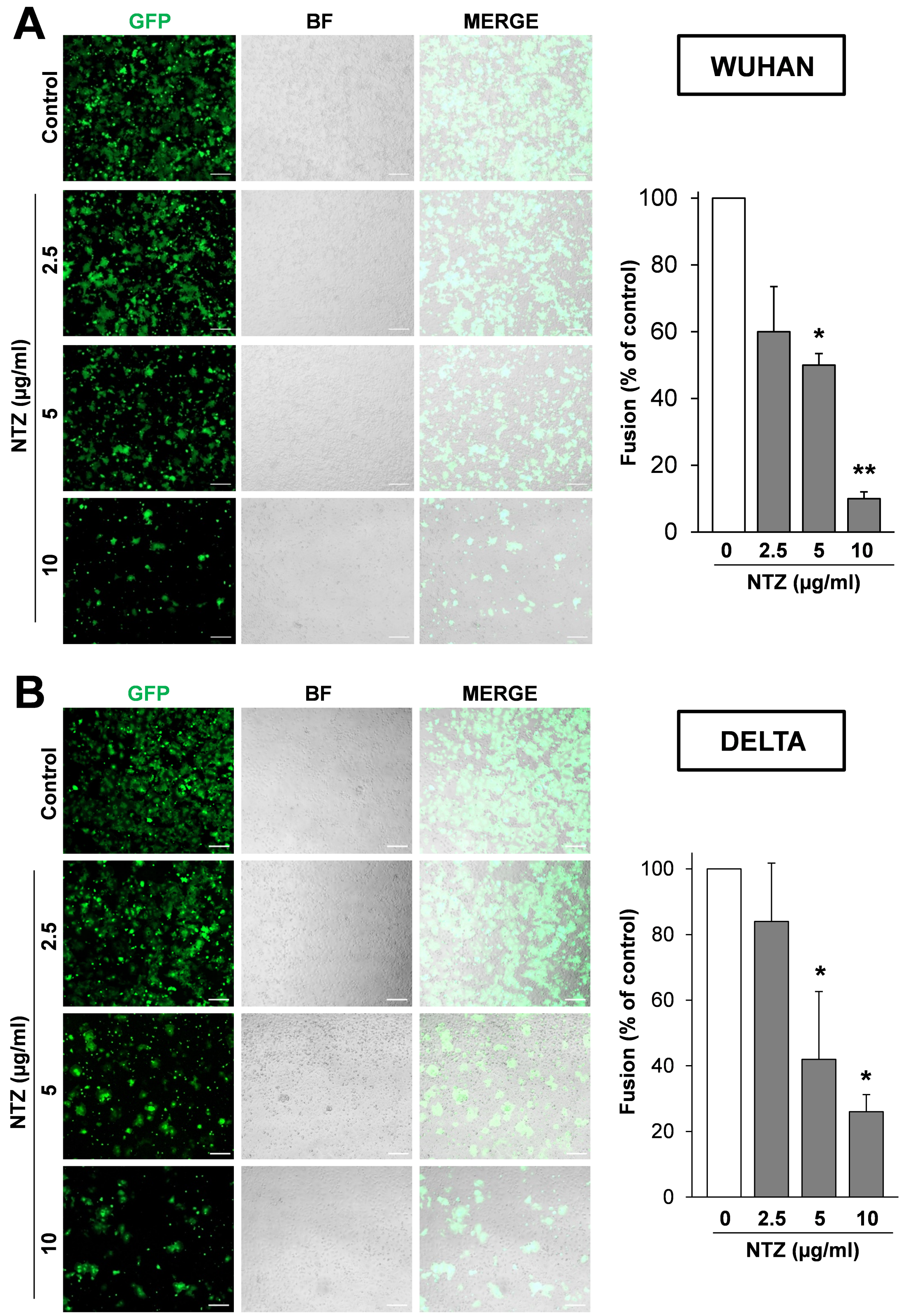


**Figure S8. Dose-dependent inhibition of SARS-CoV-2 spike-driven lung cell syncytia formation by nitazoxanide.**
